# Syntaxin1 Ser14 Phosphorylation is Required for Non-Vesicular Dopamine Release

**DOI:** 10.1101/2022.05.27.493791

**Authors:** A Shekar, SJ Mabry, MH Cheng, JI Aguilar, S Patel, D Zanella, DP Saleeby, Y Zhu, T Romanazzi, P Ulery-Reynolds, I Bahar, AM Carter, HJ Matthies, A Galli

## Abstract

Amphetamine (AMPH), a psychostimulant commonly prescribed for the treatment of neuropsychiatric and neurological disorders, has a high liability for abuse. The abuse and psychomotor stimulant properties of AMPH are primarily associated with its ability to increase dopamine (DA) neurotransmission. This increase is mediated, in large part, by non-vesicular DA release (DA efflux). DA efflux is the result of reversal of the DA transporter (DAT) promoted by AMPH. Syntaxin 1 (Stx1) is a SNARE protein that plays a pivotal role in vesicular release. Previously, we have shown that Stx1 also interacts with the distal DAT N-terminus, an event promoted by AMPH. Stx1 is phosphorylated at Ser14 by casein kinase II (CK2). Using *Drosophila Melanogaster* as an animal model, we show that this phosphorylation event is critical for non-vesicular DA release and regulates the expression of AMPH preference as well as the ability of AMPH to promote mating drive. We also show that reverse transport of DA mediated by DAT underlies these complex behaviors promoted by AMPH. Our molecular dynamics (MD) simulations of the phosphorylated DAT/Stx1 complex demonstrate that the phosphorylation state of these proteins plays a key role in allowing DAT to dwell in an efflux-willing state. This state also supports constitutive DA efflux (CDE), an event that occurs in the absence of AMPH. The DAT-Stx1 phosphorylated complex is characterized by the breakdown of two key salt bridges in DAT, K66-D345 and E428-R445, which are critical for the formation of the intracellular (IC) gate and for transport function. The breaking of these salt bridges leads to an opening and hydration of the DAT intracellular vestibule, allowing DA to bind from the cytosol, a mechanism that we hypothesize leads to CDE. We further determine the importance of Stx1 phosphorylation in CDE by pharmacologically inhibiting CK2 with CX-4945, a molecule currently in phase II clinical trials for cancer treatment. CX-4945 treatment prevented the expression of CDE in isolated *Drosophila Melanogaster* brains as well as behaviors associated with CDE. Thus, our results suggest that Stx1 phosphorylation is a possible pharmacological target for the treatment of AMPH abuse.

## Introduction

Amphetamine (AMPH) is a psychostimulant commonly used clinically to treat neuropsychiatric disorders, most notably attention deficit disorders^1^. There are estimated to be 56 million AMPH users world-wide^2^. In 2020 the Substance Abuse and Mental Health Services Administration reported that about 2.6 million people in the United States have used AMPHs in the past year while approximately 1.5 million have AMPHs use disorders, highlighting the abuse potential of AMPH^3–4^. Since 2008, hospitalizations related to AMPH and its derivatives (e.g. methamphetamine) have increased to a greater degree than hospitalizations associated with other abused substances^5^. Furthermore, in 2015, annual hospital costs associated to AMPHs abuse were $2.17 billion, underscoring the public health impact and the societal cost of these drugs^5^ as well as the need to uncover new pharmacological targets for the treatment of AMPH abuse.

The abuse potential and psychomotor stimulant properties of AMPHs have been associated with their ability to cause mobilization of cytoplasmic dopamine (DA), leading to an increase in extracellular DA levels^3, 6–7^. This increase is mediated by non-vesicular DA release (DA efflux). This non-vesicular release is the result of reversal of the DA transporter (DAT),^3^ promoted by AMPH. The physiological function of DAT is to shape central DA neurotransmission *via* re-uptake of synaptically released DA^8–9^. Notably, inhibition of DA efflux reduces both the ability of AMPH to increase motor activity and AMPH preference^10–13^. Thus, identifying the molecular underpinnings of AMPH-induced DA efflux, understanding how they impact DA neurotransmission, and translating these events to specific behavioral phenotypes associated with AMPH exposure are pivotal steps for providing new therapeutic targets for AMPH abuse.

AMPH is a substrate for DAT and is transported across the plasma membrane in a Na^+^ dependent manner, similar to DA^3, 14–15^. Uptake of a substrate through DAT is determined by five conformation states that the transporter adopts: Outward-open (O_o_), outward-occluded (O_occ_), holo-occluded, inward-occluded (I_occ_), and inward-open (I_o_)^16–17^. In the O_o_ state, the binding pocket is accessible to water and substrate, as well as sodium and chloride ions that bind in the immediate vicinity of the substrate^16^. DA, along with its cotransported ions, bind to DAT in the O_o_ conformation. This binding induces a shift through the occluded states as intermediates. Molecular dynamics (MD) simulations have facilitated additional insight into the mechanism of DA transport. Notably, it was found that in the holo-occluded state, both the extracellular (EC) and intracellular (IC) gates are closed and both vestibules are inaccessible to water^17^. In this conformation, there is formation of a salt bridge, E428-K260, and breaking of E428-R445 and K66-D345 salt bridges that control the IC vestibule gate^18^. This shift allows the transporter to begin the transition to the I_o_ state^17^. Once it reaches the I_o_ state, DA and its cotransported ions are released in to the cytosol.

In addition to acting as a DAT substrate, AMPH also induces DA efflux^3, 6, 19–21^. AMPH stimulates calcium/calmodulin-dependent protein kinase II (CAMKII), as well as protein kinase C (PKC), that in turn phosphorylate the DAT N-terminus^22–26^. This phosphorylation is required for DA efflux to occur^21,24–25^. There are 5 main phosphorylation sites at the distal N-terminus, which include Ser2, 4, 7, 12, and 13^27^. Deletion of the 22 most distal amino acids, or substitution of these Ser to Ala (S to A), do not alter DAT surface expression, surface localization, or DAT-mediated DA uptake, while it does impair AMPH-induced DA efflux^21, 27–28^. However, how phosphorylation of the DAT N-terminus promotes DA efflux and the molecular players involved in this process is still largely unknown.

Syntaxin 1A (Stx1) is a soluble N-ethylmaleimide-sensitive factor attachment protein receptor (SNARE) protein that plays a key role in vesicular synaptic release^29^ and is regulated by Munc18-1 binding^30–31^. In addition to its role in vesicular fusion, Stx1 interacts with and regulates the function of transmembrane proteins, including ion channels and transporters^12, 32–37^. Stx1 directly interacts with the DAT N-terminus within the first 33 amino acids^33^. This DAT domain includes the 5 most distal Ser whose phosphorylation is involved in DA efflux ^19, 21^. Interestingly, AMPH promotes DAT-Stx1 association as well as Stx1 phosphorylation at Ser14^12, 33^. This association occurs at the plasma membrane^33^ and is regulated by Stx1 phosphorylation, an event mediated by casein kinase 2 (CK2)^12, 38^. In addition to regulating AMPH actions^12^, the pivotal role played by Stx1 phosphorylation in neuropsychiatric disorders associated with DA dysfunction is underscored by the involvement of CK2 impairments in the etiology of both autism and schizophrenia^39^.

In this study, we delineate the temporal dynamics of AMPH-induced Stx1 phosphorylation and the role played by this phosphorylation on non-vesicular DA release and its associated behaviors. Furthermore, to understand the molecular mechanisms of how Stx1 phosphorylation promotes AMPH-induced DAT-mediated DA efflux, we performed molecular dynamics (MD) simulations. Our results support a model where AMPH, by coordinating DAT N-terminus and Stx1 phosphorylation, causes DAT to form an aqueous channel-like pore where DA molecules bind the DAT intracellular vestibule from the cytoplasm and thus cross the plasma membrane to support DA efflux.

## Materials and Methods

### DNA constructs

The pEGFP vector (ClonTech, Mountain View, CA) harboring synhDAT, synhDAT SD (hDAT with Ser2, 4, 7, 12, and 13 all mutated to Asp) and synhDAT SA (hDAT with Ser2, 4, 7, 12, and 13 all mutated to Ala) coding sequences and the pcDNA3.1 vector containing Stx1, Stx1 S14A, or Stx1 S14D sequences were generated as previously described^12^.

### Cell culture

Human Embryonic Kidney (HEK293T) cells were maintained in a 5% CO_2_ incubator at 37°C in Dulbecco’s Modified Eagle Medium (DMEM) supplemented with 10% fetal bovine serum (FBS), 1 mM L-glutamine, 100 U/mL penicillin, and 100 μg/mL streptomycin. hDAT and Stx1 constructs were transiently cotransfected into these cells (hDAT cells) using Fugene-6 (Roche Molecular Biochemicals) per the manufacturer’s protocol. Biochemical and electrophysiological assays were conducted 48-72 hours post transfection.

### [^3^H]DA Uptake Assays

Cells were plated on poly-D-lysine coated 24-well plates, transiently transfected at 24 hours post-plating, then grown to ~90% confluence. *Drosophila* brains were dissected in ice-cold Schneider’s *Drosophila* Medium (Gibco, Waltham, MA) containing 5% BSA with surgical forceps. On the day of the experiment, cells or brains were washed once with 37°C KRH buffer containing 10 mM dextrose, 100 μM pargyline, 1 mM tropolone, and 100 μM ascorbic acid, and equilibrated for 5 minutes at 37°C. Saturation kinetics of DA uptake (or single point DA uptake) was determined using a mixture of [^3^H]DA (PerkinElmer Life Sciences, Waltham, MA) and unlabeled DA diluting to final DA concentrations of 0.01 μM – 10 μM. Uptake was initiated by bath addition of each DA dilution. Uptake was terminated after 10 minutes by washing cells twice in ice-cold KRH buffer. MicroScint Scintillation fluid (PerkinElmer Life Sciences) was added to the wells and the plates were counted in a TopCount Scintillation Counter (Perkin Elmer Life Sciences). Nonspecific binding was determined in the presence of 10 μM cocaine. Km and V_max_ values were derived by fitting Michaelis-Menten kinetics to the background corrected uptake data, using GraphPad Prism 9.1 (GraphPad Software, San Diego, CA). Values reported from transport studies are derived from 3 – 5 replicate experiments.

### Cell amperometry and patch-clamp electrophysiology

Cells were washed three times with Lub’s external (130 mM NaCl, 10 mM HEPES, 34 mM dextrose, 1.5 mM CaCl_2_, 0.5 mM MgSO_4_, 1.3 mM KH_2_PO_4_, pH 7.35 and 300-310 mOsms/L) bath solution at room temperature. The internal solution for the whole-cell recording contained 120 mM KCl, 10mM NaCl 0.1 mM CaCl_2_, 2 mM MgCl_2_, 1.1 mM EGTA, 10 mM Hepes, and 30 mM dextrose plus DA (2 mM or as specified in the text) adjusted to pH 7.35. Patch electrodes were pulled from quartz pipettes on a P-2000 puller (Sutter Instruments, Novato, CA) and filled with the internal solution. Whole cell currents were recorded at a frequency of 1,000 Hz using an Axopatch 200B (Axon Instruments, Hawthorn, Australila) with a low-pass Bessel filter set at 1,000 Hz. Current-voltage relationships were generated using a voltage step (1 s) protocol ranging from −120 mV to 120 mV separated by 20 mV steps from a given holding potential.

For experiments utilizing amperometry, a carbon fiber electrode connected to a second amplifier (Axopatch 200B) was placed at the plasma membrane of the cell and held at +600 mV for all experiments. The carbon fiber electrodes (ProCFE; fiber diameter is 5μm) were obtained from Axon Instruments. For hDAT cell experiments, 10 μM AMPH was used to elicit DA efflux while 10 μM COC was used to reveal constitutive DA efflux (CDE). A sampling frequency of 100 Hz was used and the amperometric currents were low pass filtered at 10 Hz. Data were recorded and analyzed off-line using pCLAMP 9 software from Axon Instruments.

For experiments that involved treatment with CX-4945 (CK2i), cells were pretreated for 30 minutes with 100 nM CK2i (or VEH) in DMEM media. The cells were further exposed to 100 nM CK2i (or VEH) during the course of the experiment in Lub’s external solution. For experiments using hDAT conditions that display impaired uptake (e.g. hDAT SD), the DA was patch-loaded into the cells (see above). For experiments comparing hDAT conditions with normal uptake (e.g. hDAT WT), DA (1 mM for 12 minutes) was bath loaded into the cell in KRH (130 mM NaCl, 25 mM HEPES, 4.8 mM KCl, 1.2 mM KH_2_PO_4_, 1.1 mM MgSO_4_, 2.2 mM CaCl_2_, 10 mM d-glucose, pH 7.4).

### Western Blotting Protocol

Cells were washed three times with 4°C PBS (Gibco) containing 1 mM EGTA and 1 mM EDTA then lysed in RIPA buffer (100 mM NaCl, 1.0% IGEPAL CA-630 (NP-40), 0.5% sodium deoxycholate, 0.1% SDS, 50 mM Tris, 100 μM PMSF, pH = 8.0), supplemented with a protease inhibitor cocktail (Sigma Aldrich, St. Louis, MO). Lysates were passed twice through a 27.5-gauge needle and centrifuged at 15,000 x g for 30 minutes. Supernatants were then separated by SDS-PAGE gel, transferred to polyvinylidene fluoride membrane (PVDF) (Millipore, Bedford, Massachusetts) and immunoblotted. Blocking was done with either Intercept blocking buffer (Licor, Lincoln, Nebraska) or 4% milk. Primary antibodies used targeted hDAT (1:1000, MAB369 Millipore, Burlington, MA), Stx1 (1:1000, S0664 Sigma), pS14 Stx1 (1:1000, ab63571 Abcam, Cambridge, UK), and β-actin (1:1000, A2228, Sigma). Secondary antibodies were goat-anti-rat-HRP-conjugated (1:5000, Jackson ImmunoResearch, West Grove, PA). goat anti-rabbit IgG 680RD (1:10000, 926-68071, Licor), goat anti-mouse IgG 800CW (1:10000, 926-32210, Licor), or goat anti-rat IgG 800CW (1:10000, 926-32219, Licor). Band densities were quantified using ImageJ or Image Studio (Licor) and data from 3 – 4 separate experiments were combined.

### Immunoprecipitation Assays

Cells were grown and lysed in the same manner as the above methods for western blots except a Tris buffered saline (TBS), pH 7.4 with 1% Triton-X 100 (TBS-Triton) was used to lyse the cells. A portion of the total cell supernatant was collected to analyze for total protein. Remaining supernatant was incubated at 4°C for 1 hour with Sepharose-G beads (Fisher Scientific), previously washed with 5% BSA in TBS-Triton buffer and preincubated overnight with 2.5 μg anti-GFP antibody ab290 (Abcam). For the negative control, supernatant was incubated with BSA-blocked Sepharose-G beads alone (no antibody). As an additional control, lysate from mock transfected cells was incubated at 4°C for 1 hour with Sepharose-G beads (Fisher Scientific), previously washed with 5% BSA in TBS-Triton buffer and preincubated with 2.5 μg anti-GFP antibody. Beads were spun down, washed with cold RIPA buffer, and samples eluted first with sample buffer (1% SDS, 18% glycerol, 0.05% phenol blue, 62.5mM Tris HCl, 7M Urea, 2M Thiourea, 0.1M DTT) at 95°C for 5 minutes and then thiourea/urea mixture (7M Urea, 2M Thiourea) at 95°C for 5 minutes. Total lysates and eluates were analyzed by SDS-PAGE and immunoblotting (see above for antibody details). Band intensity was quantified using Image Studio (Licor). The association between Stx1 variants and hDAT variants was represented as the ratio of eluate:total lysate band intensity, normalized to the eluate:total lysate ratio observed in hDAT WT with vehicle condition and expressed as a percentage.

### Biotinylation Assays

Cells were grown and lysed in the same manner as the above methods for western blots. On the day of the assay, cells were washed three times with 4°C phosphate-buffered saline (Gibco) and sulfosuccinimidyl-2-(biotinamido)ethyl-1,3-dithiopropionate-biotin (sulfo-NHS-SS-biotin) (1.0 mg/ml, Pierce, Rockford, IL) was added to each well and allowed to incubate for 30 min at 4 °C. Cells were quenched with 100 mM glycine and then solubilized in RIPA buffer (see above for details) for 30 min at 4 °C. Extracts were collected from wells and centrifuged for 30 min at 15,000 × g at 4 °C. The supernatant was added to immunopure immobilized streptavidin beads (Pierce). Beads were allowed to incubate for 45 min at room temperature, were extensively washed, and Laemmli buffer with 2-mercaptoethanol was added. The eluent from the beads, as well as the biotinylated input (5 μg), was run on an SDS-PAGE gel, transferred to PVDF membrane (Millipore) and immunoblotted (see above for antibody details). Band densities were quantified using ImageJ or Image Studio (Licor). Surface expression was represented as surface/total band intensity and normalized to the ratio observed in hDAT WT-Stx1 group and expressed as a percentage. Data from 4 separate experiments were combined.

### *Drosophila* Rearing and Stocks

All *Drosophila melanogaster* lines were maintained on standard cornmeal-molasses fly food (Nutrifly) at 25°C under a 12:12 h light-dark schedule. Fly stocks include TH-GAL4 (Bloomington Indiana Stock Center (BI) 8848), DAT^MB07315^ (BI 25547), *w^1118^* (BI 6326), UAS-mCherry (Kyoto Stock Center 109594), M[vas-int.Dm]ZH-2A; (M[3×P3-RFP.attP’]ZH-22A (Bl 24481), and *DAT^fmn^* (dDAT KO). *Drosophila* expressing homozygous *DAT^fmn^* (dDAT KO), TH-Gal4, and UAS-mCherry were outcrossed to a control line (*w^1118^*) for 5 – 10 generations. The transgenes (hDAT WT and hDAT SD) were cloned into pBID-UASC and constructs were injected into embryos from M[vas-int.Dm]ZH-2A; (M[3×P3-RFP.attP’]ZH-22A. Flies containing transgenes were outcrossed to dDAT KO flies (in w^1118^ background) for 5–10 generations. Adult male flies (2-6 days post ecolosion) containing a single copy of hDAT WT or hDAT SD in DA neurons in the *DAT^fmn^* background were used for all behavior and electrophysiological experiments (adult female flies also used during the courtship assay; see below).

### *Drosophila* Brain Amperometry

Drosophila brains from hDAT WT or hDAT SD flies were dissected in cold Schneider’s *Drosophila* Medium (Gibco). Brains were transferred to a 35 mM sylgard dish with 3 mL Lub’s external solution (see above for details) and pinned to the plate using tungsten wire (California Fine Wire Company, Grover Beach, California). A carbon fiber electrode connected to an Axopatch 200B amplifier (Axon Instruments) was held at +600 mV was placed in to the TH-positive PPL1 DA neuron region, guided by mCherry fluorescence. For *Drosophila* brain amperometry experiments, 20 μM AMPH was used to elicit DA efflux while 1 μM COC was used to reveal CDE. The electrode, equipment, and filtering parameters were the same as utilized in the cell experiments (see above). Data were recorded and analyzed off-line using pCLAMP 9 from Axon Instruments.

### *Drosophila* Locomotion

Unmated adult male flies (2-3 days post eclosion) were collected, anesthetized with CO_2_, and transferred to glass tubes with standard fly food. *Trikinetics Drosophila* Activity Monitoring system (Waltham, Massachuesetts) was used to measure locomotion. For the circadian locomotion, after 24 hours of acclimation, locomotion was recorded for 24 hours by beam breaks. For the AMPH-induced locomotion, flies were pretreated for 24 hours with either vehicle (DMSO) or 100 nM CK2i in standard fly food. Flies were then fed standard fly food with AMPH (10 mM) and either vehicle or CK2i and, after 1 hour of acclimation, beam breaks were measured as above.

### *Drosophila* Courtship

*Drosophila* courtship was measured in a custom 3D printed chamber (33×10×10mm). *Amphetamine induced courtship:* Adult male flies (2-6 days post eclosion) were starved for 6 hours in a vial with 1% agar. Flies were then fed standard fly food containing 10 mM AMPH (or vehicle) for 20 minutes. Male flies were placed in to the chamber for 10 minutes of acclimation prior to the introduction of an adult female fly (2-6 days post eclosion). Courtship was recorded for 10 minutes (Sony, ExmoreR OpticalSteadyShot) *CK2i courtship:* Adult male flies were fed standard fly food containing 100 nM CK2i (or vehicle) for 24 hours prior to courtship. Male flies were placed in to the 3D printed chamber and the same acclimation and recording protocol as above was performed.

All courtship videos were scored blinded to the genotype and drug treatment. Video scorers manually timed and recorded the time spent displaying one of five courtship behaviors (following, wing extension, abdomen bend, tapping, and attempted copulation). The courtship index was calculated as the fraction of the amount of time the male spent courting the female over the total 10 minutes. Latency to court was measured as duration of time between introduction of the female and displaying of courtship behavior by the male. Achievement of copulation was recorded and utilized for analysis.

### *Drosophila* Two Choice Assays

A custom two choice apparatus was constructed to measure drug preference at the nL level in *Drosophila.* Each apparatus contained two volumetric capillaries: one with clear food (100 mM sucrose) and one with blue food (100 mM sucrose, 500 μM blue dye). Half of the groups also received CK2i (100 nM) in both capillaries while half of the groups received VEH (DMSO) in both capillaries. Food consumption was measured every 24-hours and capillaries were refilled and replaced. Adult male flies (2-5 days post eclosion) were anesthetized by CO_2_, transferred to the apparatus, and given 24 hours to acclimate to the liquid capillary food. The following 24 hours was used to calculate baseline preference of clear and blue food. To determine AMPH preference, on the third day, the blue capillary was supplemented with 1 mM AMPH (see Figure 7A). Preference was calculated as consumption of blue food over total consumption (blue food and clear food together).

### Molecular modeling of Stx1 and Stx1-CK2 binary complex

Human syntaxin 1A (Stx1; Q16623) structural models (residues 1-260) were constructed using Robetta^40^, based on Stx1 in a closed conformation^41^ (PDB: 4JEH); and the missing loop (residues 1 to 26) in Stx1 was modeled by Robetta. Using the top five Stx1 models, we performed protein-protein docking simulations for each Stx1 model against the hCK2 (V11-V328) crystal structure (PDB: 3U4U)^42^ using ClusPro^43^. We selected one Stx1 model for further study, in which the hCK2 catalytic residue D156 is in proximity of Stx1 S14 (known phosphorylation site^38^; see Supp. Fig. 1A)

### Modeling of hDAT-Stx1 complex structure

The full length hDAT in an *occluded* conformation was taken from previous study^11, 44^. We generated a series of structural models for the hDAT-Stx1 binary complex using ClusPro^43^. We used *occluded*/closed states of hDAT and Stx1 as conformers. 1,000 models were generated that were grouped in 30 clusters based on their structural similarities. These were rank-ordered based on cluster size (accounting for favorable entropic effects) and further sorted following three criteria specific to the hDAT-Stx1 complex in a lipid environment: (1) Stx1 should bind from the intracellular (IC) region (i.e. extracellularly (EC) – bound forms were excluded); (2) the hDAT-Stx1 complex should have proper orientation of the aromatic residues (i.e., Trp) on the EC and IC sides to enable suitable anchoring into the lipids; and (3) residues (e.g. Stx1 R26 and hDAT R51), whose mutations have been shown to alter hDAT-Stx1 interactions in previous studies^12^, should be involved directly or indirectly in interfacial contacts. Notably, the most energetically favorable model, predicted by ClusPro, satisfied these three criteria, and thus was selected for further investigation using molecular dynamics (MD) simulations (see Fig. 1D).

**Figure 1:**
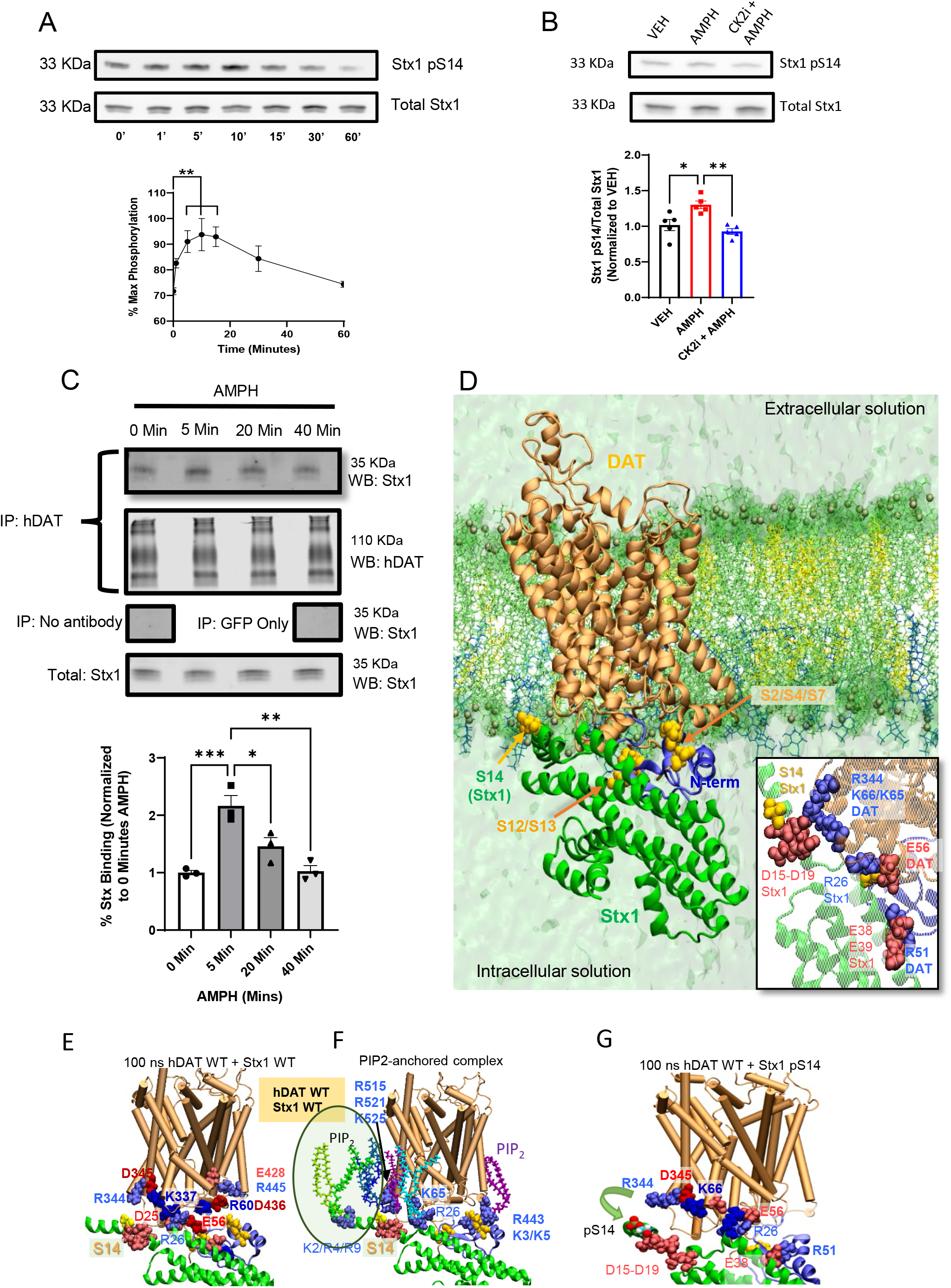
AMPH Promotes Phosphorylation of Stx1. **A.** Representative immunoblot (top) and quantitative analysis (bottom) of hDAT cells treated with 10μM AMPH for the times indicated. AMPH significantly increased Stx1 phosphorylation (F(6, 21) = 5.293, *p* = 0.0018, *n* = 4). **B.** Representative immunoblot (top) and quantitative analysis (bottom) of hDAT cells, pretreated with either vehicle or 100nM CX-4945 (CK2i) for 6 hours, then exposed to vehicle alone (VEH, black bar) or 10μM AMPH for 10 minutes in VEH (red bar) or CK2i (blue bar). AMPH significantly increased Stx1 phosphorylation and pretreatment with CK2i blocks this increase (F(2, 12) = 11.06, *p* = 0.0019, *n* = 5;). **C.** Representative immunoblot (top) for Stx1 and quantitative analysis (bottom) of hDAT immunoprecipitates (IP) from hDAT cells treated with 10μM AMPH for the times indicated. Data are expressed as IP Stx1/IP hDAT normalized to IP Stx1/IP hDAT at 0 minutes AMPH. AMPH significantly increased hDAT-Stx1 interaction at the 5 and 20-minute time points (F(3, 8) = 17.62, *n* = 3, *p* = 0.0007). **D.** MD simulations of the hDAT/Stx1 complex embedded into neuronal lipids. *Red* and *blue* Van der Walls (VDW) spheres represent acidic and basic residues. Residues from DAT and Stx1 are labeled in bold-face and plain-face, respectively. The hDAT/Stx1 complex model predicted by *ClusPro* (DAT shown in *orange* with the N-terminus shown in *blue)* was embedded into neuronal lipids *(licorice)* containing POPE and POPC (*green*), cholesterol *(yellow),* POPS (*cyan*), and PIP_2_ *(blue).* This snapshot is at 10 ns equilibration. S14 of Stx1 and the five most distal N-terminus serines (S2, S4, S7, S12 and S13) of DAT are shown in *orange* VDW spheres. *(inset)* Predicted interfacial contacts for hDAT and Stx1. **E.** Formation of IC salt bridges R60-D436, R445-E428 and K66-D345 at 100 ns equilibration, indicating DAT remained in the occluded conformation. Note that the predicted interfacial salt bridges remained, e.g. Stx1 E25-hDAT K337, Stx1 R26-hDAT E56, Stx1 E38-hDAT R51 (shown in *inset).* **F.** PIP_2_ binding coordinates some of the interaction of hDAT with Stx1. PIP_2_ lipids are colored differently depending on the site of interaction and displayed in the licorice format. **G.** MD simulations at 100ns of hDAT and Stx1 pS14. Phosphorylation of Stx1 S14 destabilized the segment pS14-D19 and promotes its dissociation from hDAT while the Stx1 R26-hDAT E56 salt bridge was retained (see Supp. Fig. 1 for entire simulation). Data are presented as mean ± SEM. One-way ANOVA with Dunnett’s multiple comparisons test (A) or Tukey’s multiple comparisons test (B, C).

### Molecular Dynamics (MD) simulations of hDAT-Stx1 complex in the neuronal lipid environment

Three simulation systems were constructed using CHARMM-GUI Membrane Builder module^45^, based on ClusPro-predicted binary complex model, and simulations of 100 ns each were conducted in triplicate for each system:

1. hDAT SD + Stx1 S14A,
2. hDAT WT + Stx1 pS14, in which Stx1 S14 was phosphorylated; and
3. hDAT SD + Stx1 pS14, in which Stx1 S14 was phosphorylated.
4. hDAT WT + Stx1 WT

For each system, the protein complex was embedded into neuronal membrane/lipids composed of 1-palmitoyl-oleoyl-sn-glycero-3-phosphoethanolamine (POPE) and -phosphocholine (POPC), cholesterol (CHOL), palmitoyl-oleoyl-phosphatidylinositol (POPI), and phosphatidylinositol 4,5-bisphosphate (PIP_2_) using CHARMM-GUI Membrane Builder module^45^. The lipid composition was adapted from previous study^46^ to account for the asymmetric lipid distribution of neuronal membranes: the outer lipid layer contained POPC (35%), POPE (35%), and CHOL (30%); and the inner contained POPC (20%), POPE (25%), CHOL (25%), PIP_2_ (15%), and POPS (15%) with the percentage of lipids in each layer written in parentheses. Fully equilibrated transferable intermolecular potential 3P (TIP3P) waters were added to build a simulation box of 136×136 ×167 Å^3^. Na^+^ and Cl^-^ ions were added to obtain a 0.15M NaCl neutral solution. Each simulation system contained approximately 291,000 atoms, including those of the hDAT-Stx1 complex, 570 lipid molecules, and 70,000 water molecules. A typical MD simulation setup is shown in Fig. 1D. For the system (3), i.e. hDAT SD + Stx1 pS14, ten dopamine molecules were distributed to both EC and IC solutions. All simulations were performed using NAMD^47^ following previous protocol^46^. The probability of selected interactions was assessed by counting the times of atom-atom interactions within 4 Å distance for the examined amino acid pair, at 0.2 ns intervals during 100 ns trajectories. VMD^48^ with in-house scripts was used for visualization and trajectory analysis.

### Statistical Analyses

Experiments were designed after statistical power calculations based on preliminary data using GPower 3.1. Statistical analyses were run on GraphPad Prism 9.1 (San Diego, CA). Shapiro-Wilk normality tests were performed to ensure data was normally distributed and Bartlett’s tests were performed to check for heteroscedasticity. If data failed these assumptions the proper non-parametric tests were utilized. Unpaired t-tests were used for all comparisons between two groups and Welch’s correction was used whenever the variances significantly differed between groups. When using one-way or two-way ANOVA’s, Tukey’s post hoc was used in most cases. However, in the scenario that all means were being compared to a control mean, Dunnett’s post hoc was used. In the case that the post hoc comparisons were only done between the different concentrations between groups, such is the case in the kinetic uptake experiments, Bonferonni’s post hoc was used. For all repeated measures two-way ANOVA’s, Sidak’s multiple comparisons post hoc was used. For categorical data, Chi-square test was used. Mantel-Cox Log-Rank survival analysis was used for all survival plots. For all experiments * = *p* < 0.05, ** = *p* < 0.01, *** = *p* < 0.001, **** = *p* < 0.0001.

## Results

### Amphetamine Promotes Stx1 Phosphorylation and Reorganizes the hDAT/Stx1 Complex

AMPH induces CK2-mediated Stx1 phosphorylation at S14^12^ and binding of Stx1 to the DAT N-terminus^12, 33, 49^. These events are vital for the ability of AMPH to cause DA efflux and associated behaviors^12, 33^. We defined the temporal dynamics of AMPH-induced Stx1 phosphorylation, the key players implicated, as well as the structural domains involved in DAT-Stx1 interactions. In hDAT cells (see methods), AMPH treatment causes a significant increase in Stx1 phosphorylation at S14 (pS14) at 5, 10, and 20 minutes. However, after 30 minutes of AMPH exposure, the level of Stx1 phosphorylation was not significantly different with respect to baseline (Fig. 1A). To further explore this AMPH-induced phosphorylation event, we focused on CK2, a kinase implicated in phosphorylation of Stx1 at S14^12, 38^. We treated hDAT cells with the CK2 inhibitor CX-4945 (CK2i)^50^ before (6 hours) and during AMPH exposure. We found that CK2i blocks Stx1 phosphorylation in response to AMPH treatment (Fig. 1B). Consistent with this CK2 activity, docking simulations between CK2 and Stx1 support a model where CK2 is capable of phosphorylating Stx1 at S14 (Supp. Fig. 1A), i.e. Stx1 S14 makes close contact with the CK2 catalytic residue D156. These data demonstrate that CK2 mediates AMPH-induced Stx1 phosphorylation, this phosphorylation is time dependent, and that it peaks shortly after AMPH administration.

Considering that AMPH-induced hDAT-Stx1 interaction has been shown to regulate DA efflux^33^ and behaviors^12^, we next defined the temporal dynamics of AMPH induced hDAT-Stx1 associations. hDAT cells were exposed to AMPH for different time periods and hDAT was immunoprecipitated. The immunoprecipitates were then immunoblotted for Stx1 (Fig. 1C; top). The interaction between hDAT and Stx1 peaked at 5 minutes of AMPH exposure (Fig. 1C; bottom), a time that closely corresponds to that of maximum Stx1 phosphorylation (Fig. 1A). Thus, it is possible that the interactions between hDAT and Stx1 are tightly regulated by Stx1 phosphorylation status. Consistent with this hypothesis, this interaction returned to baseline by 40 minutes (Fig. 1C; bottom) similar to what observed for Stx1 phosphorylation (Fig. 1A).

The question remains to understand how Stx1 phosphorylation at S14 regulates the structural interactions between hDAT and Stx1. First, we generated structural models for the complex of hDAT WT with Stx1 WT using full length hDAT in an *occluded* conformation^44^ and the Stx1 structure modeled by Robetta^51^, using ClusPro^43^. Then we performed MD simulations of hDAT-Stx1 complex in the presence or absence of a phospho group at Stx1 S14 (Stx1 pS14). In Figure 1D, we show the resulting complex formed by hDAT and Stx1 embedded in a membrane^46^ with surrounding water molecules, ions, and lipids including PIP2, obtained after 10 nanoseconds MD simulations. In this complex, the interfacial interactions between hDAT and Stx1 involve multiple salt bridges: Stx1 D15/D17 and hDAT R344, Stx1 R26 and hDAT E56, and Stx1 E38/E39 and hDAT R51 (Fig. 1D, inset; Fig. 1E; 100 ns snapshot). Notably, mutations associated with autism (R26Q in Stx1 and R51W in hDAT), which would result in disruption of these predicted salt bridges, have been implicated in altering hDAT-Stx1 interactions, Stx1 phosphorylation, as well as DA efflux^12^. The N-terminal segment of Stx1 (M1-R26) is highly charged, carrying four positive and eight negative charges. Five contiguous acidic residues (D15-D19) form a negative cluster in proximity to S14. In contrast, the cytoplasmic side of hDAT contains a cluster of basic residues that attract the acidic portions of Stx1. In particular, the basic cluster in hDAT containing R344 and K65/K66 is predicted to be near the D15-D19 Stx1 acidic cluster (Fig. 1D, inset).

We also found that PIP_2_ coordinates some of the hDAT WT-Stx1 interactions. In our MD simulations, we consistently observed two to four PIP_2_ molecules binding to the interface between hDAT WT (R515, R521 and K525) and the N-terminus of Stx1 (K2, R4 and R9) (circled in Fig. 1F), which is consistent with the observation that Stx1 N-terminus interacts with PIP_2_^52^. The hDAT WT/Stx1 complex includes two additional PIP_2_ binding sites: one *(cyan licorice,* Fig. 1F) is near K65/K66 of DAT and R26/R28 of Stx1, and the other *(purple licorice,* Fig. 1F) is near K3/K5 and R443 of DAT. Consistent with this model, we previously identified a basic cluster (including K3 and K5) at the N-terminus and the fourth intracellular loop (IL4; R443) as PIP_2_ binding sites^10–11^. In the MD simulations of the complex between hDAT WT and Stx1 pS14 complex, the phosphorylation of S14 destabilized the S14-D19 segment of Stx1 and triggered its dissociation from hDAT WT K65/K66 and R344 (Fig. 1G, snapshot at 100ns; Supp. Fig. 1B-D).

### Phosphorylation of Stx1 at Ser14 Supports AMPH-Induced DA Efflux

AMPH’s ability to cause DA efflux dictates its rewarding and addictive properties^11, 13^. Cleaving Stx1 or preventing Stx1 phosphorylation reduces AMPH-induced DA efflux as well as AMPH-induced locomotion^12^. Thus it is pivotal to understand the role played by Stx1 phosphorylation at S14 in the reverse transport of DA. We manipulated Stx1 phosphorylation both genetically and pharmacologically and measured DA efflux by amperometry in hDAT cells. AMPH-induced DA efflux (represented as an upward deflection of the amperometric trace) was significantly reduced by inhibiting Stx1 phosphorylation with CK2i (Fig. 2A). Furthermore, hDAT cells expressing Stx1 S14A (prevents S14 phosphorylation) exhibited significantly reduced efflux when compared to hDAT cells expressing Stx1 (Fig. 2B). In contrast to what was observed in cells expressing Stx1, CK2 inhibition failed to reduce DA efflux in cells expressing Stx1 S14D (Asp behaves as a phosphomimetic) (Fig. 2C). These data, combined with the data in Fig. 1B, demonstrate that AMPH-induced CK2-mediated Stx1 phosphorylation at S14 is critical for AMPH-induced DA efflux.

**Figure 2:**
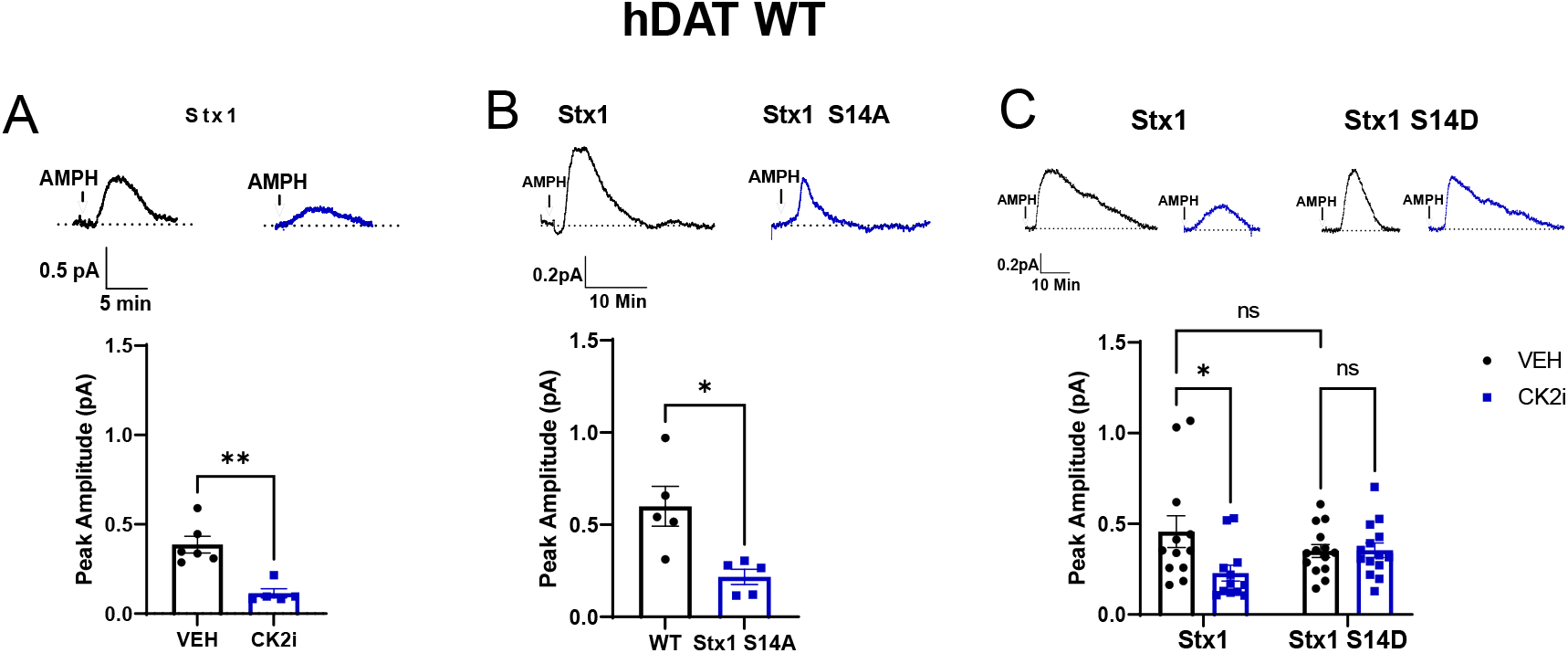
Stx1 Phosphorylation Supports AMPH-induced DA Efflux. **A.** (*top*) Representative traces of AMPH-induced DA efflux in hDAT cells treated with CK2i (100nM for 30 min) or VEH. *(bottom)* quantitative analysis of peak amplitude of the amperometric currents. CK2i treatment significantly reduced DA efflux (U = 0, *n* = 5-6).**B.** Representative traces (top) and quantitative analysis (bottom) of hDAT WT cells expressing either WT or Stx1 S14A. Stx1 S14A demonstrates a diminished ability to support DA efflux (t = 3.307, *n* = 5). **C.** Representative traces (top) and quantitative analysis (bottom) of hDAT WT cells expressing either WT or Stx1 S14D. CK2i (100nM) treatment significantly reduced DA efflux in cells expressing Stx1 but had no effect on cells expressing Stx1 S14D (F(1, 48) = 4.756 for effect of interaction, *p* = 0.0341, *n* = 12-14). Data is presented as mean ± SEM. Mann-Whitney test (A); Student’s unpaired t-test (B); Two-way ANOVA with Tukey’s multiple comparisons test (C)

### Phosphorylation of hDAT N-terminus and Stx1 S14 Promotes Constitutive DA efflux and the Formation of a DAT Aqueous Pore in an Interdependent Fashion

AMPH causes Stx1 S14 phosphorylation (Fig. 1A) an event required for proper DA efflux (Figure 2)^12, 33^. AMPH also promotes DAT phosphorylation at the most distal N-terminal Ser^22–26^. Phosphorylation of these Ser residues is required for AMPH to promote a robust DA efflux and associated behaviors^21, 24–25^. However, it is unclear whether these phosphorylation events regulate each other and coordinate DA efflux in an interdependent fashion by regulating the structural/functional conformations of the hDAT/Stx1 complex. To answer this question, we first utilized a combination of hDAT WT, a phospho-deficient N-terminal mutant (hDAT SA; S2A, S4A, S7A, S12A, S13A) as well as a phospho-mimetic N-terminal mutant of hDAT (hDAT SD; S2D, S4D, S7D, S12D, S13D). First, we determined whether the phosphorylation status of the hDAT N-terminus dictates the ability of AMPH to cause Stx1 phosphorylation at S14. In cells transfected with hDAT WT and Stx1, phosphorylation of Stx1 S14 is elevated upon AMPH exposure (Fig. 3A and Fig. 1A). However, in hDAT SA cells AMPH fails to increase Stx1 phosphorylation (Fig. 3A). Quantitation of the immunoblots is shown in Figure 3A (bottom). These data indicate that phosphorylation of the hDAT N-terminus is required for AMPH-induced Stx1 phosphorylation. Consistent with this notion, hDAT SD cells exhibit increased level of Stx1 pS14 under basal conditions (Supp. Fig. 2A; top). This increased level of Stx1 phosphorylation is not augmented by AMPH exposure (Supp. Fig. 2A; top). Quantitation of the immunoblots demonstrates a significant difference in Stx1 pS14 levels between cells expressing either hDAT WT or hDAT SD (Supp. Fig. 2A; bottom).

**Figure 3:**
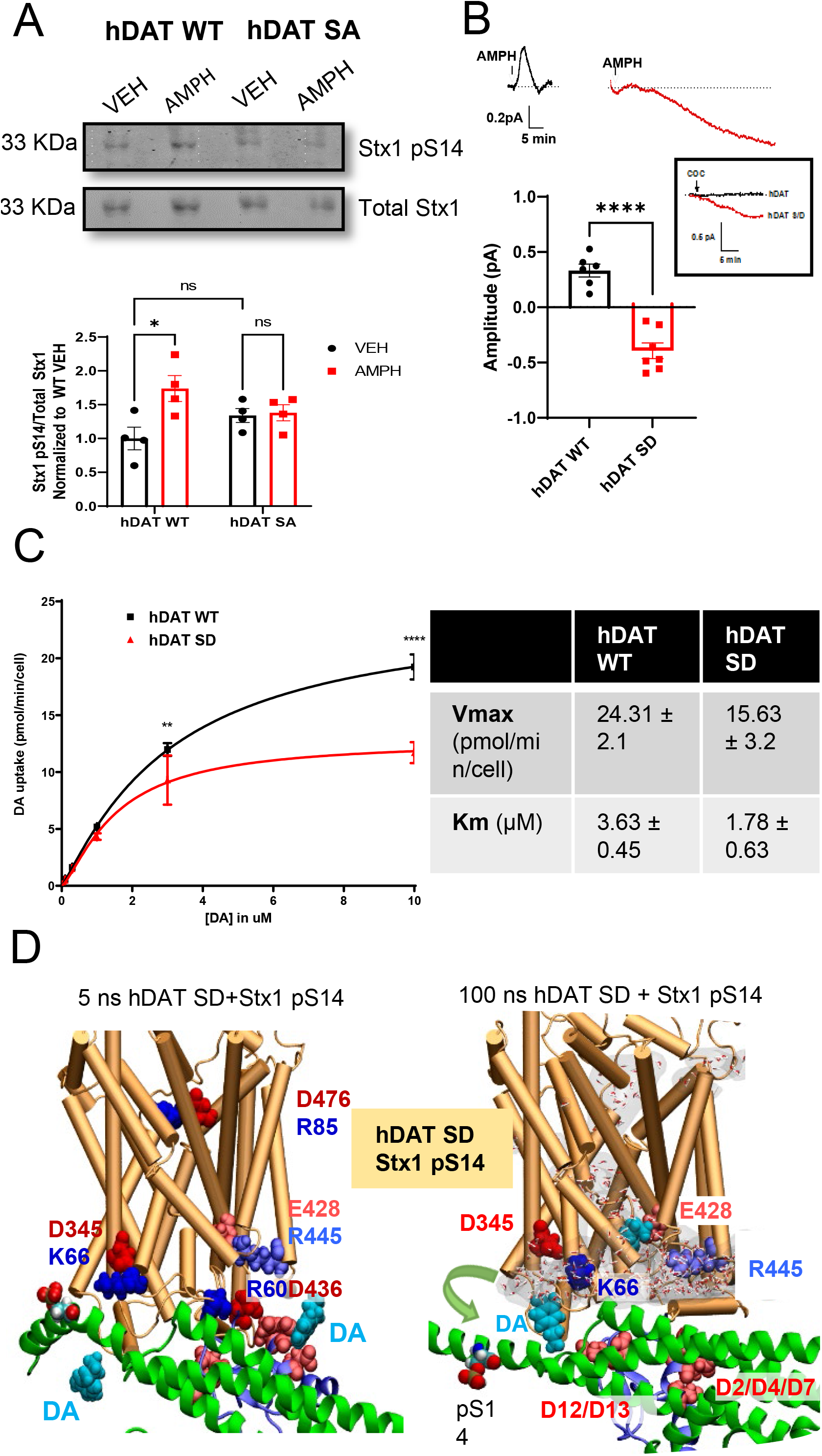
hDAT N-Terminus Pseudophosphorylation Supports Constitutive DA Efflux and the Formation of an hDAT Aqueous Pore. **A.** Representative immunoblot (top) and quantitative analysis (bottom) of Stx1 S14 phosphorylation in hDAT WT (WT) or hDAT SA (SA) cells treated with 10μM AMPH for 10 minutes. AMPH significantly increased Stx1 S14 phosphorylation in hDAT WT and failed to increase Stx1 phosphorylation in hDAT SA (F(1, 12) = 0.016 for effect of genotype, *p* = 0.951; F(1,12) = 28.04 for effect of AMPH, *p* = 0.023; F(1,12) = 22.44 for effect of interaction, *p* = 0.038, *n* = 4). **B.** Representative amperometric traces (top) and quantitative analysis (bottom) of AMPH treatment (10μM) in hDAT WT or hDAT SD cells. AMPH causes DA efflux in hDAT WT but reveals CDE in hDAT SD cells (t = 7.720, *n* = 6-7). (Inset) Representative trace (*n* = 5) of hDAT WT or hDAT SD cells treated with COC (10μM). **C.** ^3^[H]DA uptake saturation curves measured from hDAT WT (black) or hDAT SD (red) cells. Curves were fitted to Michaelis-Menten equation to derive K_m_ and V_max_. At multiple DA concentrations DA uptake for hDAT SD cells was significantly reduced compared to hDAT WT cells (F(5, 24) = 7.56, *n* = 4 in triplicate, *p* = 0.0002). V_max_ was significantly reduced in hDAT SD (*t* = 2.458, *n* = 4, *p* < 0.05) while K_m_ was not significantly changed (*t* = 1.181, *n* = 4, *p* = 0.282). **D.** Snapshots of MD simulation of the complex between hDAT SD and Stx1 pS14 complex at 5 ns and 100 ns. *Red* and *blue* vDW spheres represent acidic and basic residues. Residues from DAT and Stx1 are labeled in bold-face and plain-face, respectively. The phosphorylation of Stx1 S14 (pS14) and pseudophosphorylation of hDAT (SD) promotes the dissociation of the hDAT K66-D345 salt bridge and causes the reconfiguration of hDAT into an open, inward-facing-like conformer. DA is represented as magenta and H_2_O is shown as a ball and stick model. More details are in Supp. Fig. 4. Data is presented as mean ± SEM. Two-way ANOVA with Tukey’s Multiple Comparisons Test (A); Student’s unpaired t-test (B, C); Two-way ANOVA with Bonferroni’s multiple comparisons test (C).

We discovered that in hDAT SD cells, AMPH, rather than causing DA efflux (as observed in hDAT WT cells), initiates a downward deflection of the amperometric trace (Fig. 3B, top). This downward deflection of the amperometric current caused by AMPH, defined as constitutive DA efflux (CDE), has been previously observed in cells expressing hDAT genetic variations associated with either autism or ADHD^53–55^. The amplitude of the hDAT WT peak current as well as hDAT SD-mediated CDE (the amplitude of CDE is defined as the maximum deflection of the amperometric current with respect to baseline) is represented in Figure 3B (bottom). The continuous nature of CDE promoted by the hDAT SD mutant was also confirmed by its cocaine (COC) sensitivity (Fig. 3B, inset). COC is a known DAT antagonist and previously shown to block CDE^53, 55^. These data demonstrate that in hDAT SD cells, AMPH does not promotes DA efflux, but rather is now acting as a blocker of CDE, similarly to what is observed with COC.

Next, we further characterized the functional impact of hDAT N-terminus. We performed radiolabeled [H^3^]DA uptake in both hDAT WT and hDAT SD cells (Fig. 3C, top). While the K_m_ was not different between the two genotypes (p > 0.05), V_max_ was significantly reduced in hDAT SD cells (p < 0.05) (Fig. 3C; bottom). These results demonstrate that hDAT SD cells have reduced [H^3^]DA uptake capacity, a phenomenon we attribute to their ability to constitutively efflux DA, which would reduce intracellular accumulation of DA. To further support this notion, we found that the decreased [H^3^]DA uptake capacity does not stem from a protein trafficking phenomenon, since hDAT SD cell surface expression is comparable to that observed for hDAT WT (Supp. Fig. 3). Moreover, among the different combinations of hDAT/Stx1 mutants tested only hDAT SA/Stx1 S14A displays a membrane expression that is significantly different from hDAT WT/Stx1 (Supp. Fig. 3).

To understand, at the molecular level, how combined phosphorylation of hDAT N-terminus and Stx1 leads to CDE, we performed MD simulations of the complex formed between hDAT SD and Stx1 pS14. Figure 3D displays snapshots at 5 ns and 100 ns. In Fig. 1D (inset) we show that the D_25_RDR_28_ stretch of Stx1, which includes R26, contributes to stabilizing the association of Stx1 with hDAT through its interaction with hDAT E56. The strength of this interaction is enhanced by S12 and S13 pseudophosphorylation (S12D and S13D) at the hDAT N-terminus. Furthermore, S12D and S13D promotes the formation of a salt bridge with Stx1 R26 (Supp. Fig. 4) and may contribute to the experimentally observed increase in the affinity of hDAT to bind to Stx1 when exposed to AMPH (Fig. 1C). In all hDAT/Stx1 complexes tested, Stx1 D25, R26, D27 and R28 were observed to form intermittent salt bridges with hDAT K65/K337, E56/E61, K65/K337, and E61/D68, respectively. However, pseudophosphorylation of hDAT N-terminus and its increased interaction with Stx1 R26 promote the breaking of hDAT intramolecular salt bridges K66-D345 and E428-R445 (Fig. 3D *right* and Supp. Fig. 4) and lead to an inward-facing conformer. In the initial configuration, hDAT was in an *occluded* state, in which the EC gate pair R85-D476 and three IC gating pairs R60-D436, K66-D345 and E428-R445^17–18^ were closed (Fig. 3D *left).* During the simulations with hDAT SD and Stx1 pS14, these three IC salt bridges broke intermittently, leading to the opening of intracellular vestibule and the influx of water molecules (Fig. 3D *right* and Supp. Fig. 4A-C). Notably, one of the two displayed DA molecules (both shown in cyan *space-filling),* initially 25Å away from hDAT E428 (Fig. 3D; *left*), migrated from the cytoplasmic solution and bound inside the intracellular vestibule near hDAT E428 (Fig. 3D; *right);* and the second diffused to bind near the intracellular entrance close to K66 (Fig. 3D; *right).* The opening probabilities of the EC gate R85-D476 and the IC gate R60-D436 were similar in all hDAT/Stx1 complexes simulated, with some of these salt bridges coordinated by PIP_2_ molecules (Fig. 1F and Supp. Fig. 5). However, hDAT SD, in the presence of Stx1 pS14, decreased the probability of formation of the hDAT K66-D345 salt bridge from 54±20% to 17±15%, promoting the opening of hDAT IC gate. Thus, it is possible that the opening of the IC gate and the subsequent binding of DA from the cytosol near E428 and K66 is key for hDAT to transition from an uptake ready to an efflux ready conformation and the expression of CDE. Thus, this channel-like state of the hDAT seems to be required for CDE and is dictated by the coordinated phosphorylation of both hDAT and Stx1.

### Stx1 Phosphorylation is Required for CDE

To further understand the role of Stx1 phosphorylation in CDE, we recorded AMPH-induced DA efflux in hDAT SD cells exposed to either vehicle or CK2i (Fig. 4A). It is important to note that hDAT SD promotes Stx1 phosphorylation (Supp. Fig. 2A) and this offers the opportunity to determine the role played by Stx1 pS14 in AMPH induced DA efflux once hDAT N-terminus is phosphorylated. As already described in Fig. 3B, we found that, in the presence of VEH, AMPH inhibits CDE. However, in the presence of CK2i, AMPH causes a DA efflux that is comparable to what is observed in hDAT WT cells in the presence of CK2i (compare Fig. 4A to Fig. 2A). Thus, these diverse outcomes of AMPH treatment might be attributed to the different hDAT-Stx1 interactions that depend on the Stx1 phosphorylation status. Consistent with this notion, expression of Stx1 S14A in hDAT SD cells also restored the ability of AMPH to cause DA efflux (Fig. 4B). As expected, in hDAT SD cells expression of Stx1 S14D promotes CDE, as revealed by AMPH blockade of the amperometric signal (Fig. 4B). These data indicate that upon phosphorylation of the hDAT N-terminus, phosphorylation of Stx1 is necessary for the expression of CDE, a phenomenon possibly supported by the formation of a hDAT channellike state (Fig. 3D).

**Figure 4:**
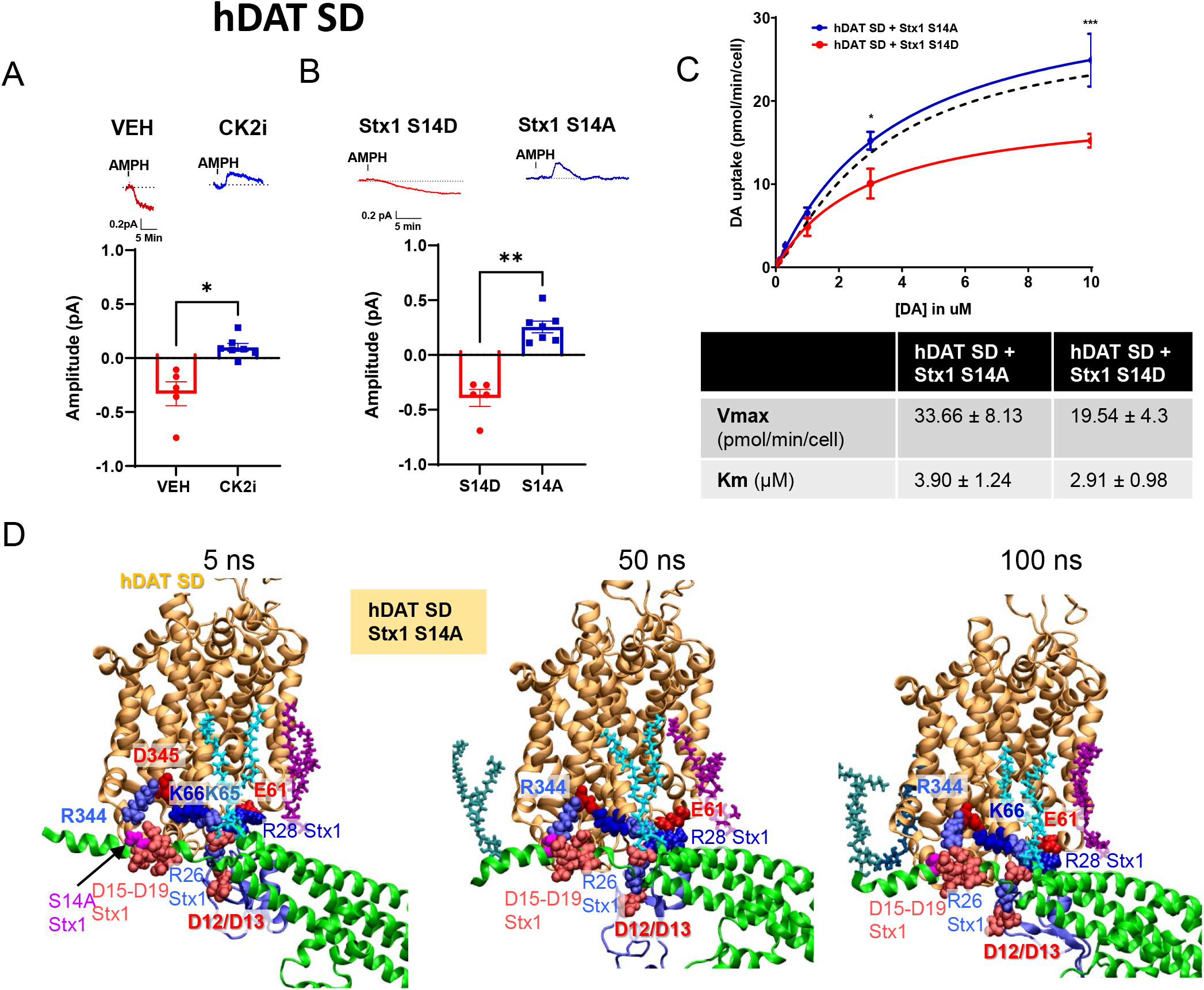
Stx1 Phosphorylation at Ser14 is Critical for CDE. **A.** Representative traces (top) and quantitative analysis of the amplitude of the peak current and CDE (the amplitude of CDE is defined as the maximum deflection of the amperometric current with respect to baseline) (bottom) of hDAT SD cells treated with CK2i (100nM) or VEH. CK2i treatment prevented the constitutive efflux (t = 3.678, *n* = 5-7). **B.** Representative traces (top) and quantitative analysis (bottom) of hDAT SD cells expressing either Stx1 S14D or S14A. Stx1 S14A prevented CDE (U = 0, *n* = 5-7). **C.** Saturation curves of ^3^[H]DA uptake measured in hDAT SD cells expressing either Stx1 S14D (red) or Stx1 S14A (blue). Curves were fit to Michaelis-Menten kinetics to derive K_m_ and V_m_a×. DA uptake for hDAT SD cells expressing Stx1 S14A was significantly increased compared to hDAT SD cells expressing Stx1 S14D cells at the two highest DA concentrations used (F(5, 22) = 4.197, *n* = 3 in triplicate, p = 0.0079). Both V_max_ (*t* = 2.659, *n* = 4, *p* = 0.056) and Km (*t* = 1.085, *n* = 4, *p* = 0.339) were not significantly different. The black dotted line represents hDAT WT cell uptake. **D.** Representative snapshots of MD simulations of the complex between hDAT SD and Stx1 S14A at 20 ns; 50 ns and 100 ns. *Red* and *blue* vDW spheres represent acidic and basic residues. Residues from DAT and Stx1 are labeled in bold-face and plainface, respectively. Note that Stx1 D25, R26, D27 and R28 were observed to form intermittent salt bridges with hDAT K65/K337, D12/D13, K65/K337, and E61/D68, respectively. For clarification, some residues, e.g. Stx1 D25, hDAT D68 and K337 are not displayed. In two independent runs, we observed similar trends, confirming the stability of the complex. In each figure, the binding PIP_2_ lipids are colored differently in *licorice* format. Note that the absence of bound PIP_2_ near R344 and K66 holds the association of D15-D19 acidic cluster of Stx1 with a basic cluster of hDAT near R344/K66. Student’s unpaired t-test with Welch’s correction (A); Mann-Whitney Test (B), Student’s unpaired t-test (C); Two-way ANOVA with Bonferroni’s multiple comparisons test (C).

We hypothesized that the reduced DA uptake observed in hDAT SD cells (Fig. 3D) was the result of CDE. Thus, inhibition of Stx1 phosphorylation, which prevents CDE, should restore normal DA uptake in hDAT SD cells. Expression of Stx1 S14A rescues the impaired [H^3^]DA uptake observed in hDAT SD cells (Fig. 4C, blue line) to that of hDAT WT (Fig. 4C, black dotted line) levels. Consistent with our hypothesis, expression of Stx1 S14D impaired DA uptake in hDAT SD cells (Fig. 4C, red line). These data demonstrate that Stx1 Ser14 phosphorylation plays a key regulatory role in determining the functional modalities of the phosphorylated hDAT conformer (uptake versus efflux).

It is important to note that in cells expressing the phosphodeficient hDAT mutant, hDAT SA, the presence of Stx1 S14D does not promote CDE (Supp. Fig. 6). These data demonstrate that phosphorylation of neither hDAT nor Stx1 alone is sufficient for the expression of CDE, but rather, combined phosphorylation of these proteins is required for CDE to occur. Considering the results of Fig. 2 and Supp. Fig. 2, they also suggest that while hDAT SD can promote Stx1 phosphorylation and CDE, Stx1 S14D is not capable of reciprocally promoting hDAT phosphorylation.

To further understand how Stx1 phosphorylation determines the different abilities of hDAT SD to respond to AMPH, we performed 100 ns MD simulations of the complex formed between hDAT SD and Stx1 S14A mutant (Fig. 4D, shown are three snapshots) and compared them to those performed for hDAT SD complexed with Stx1 pS14. Stx1 S14A substitution withstood the electrostatic interaction between hDAT SD R344 and the acidic cluster of Stx1 D15-D19 (Fig. 4D). The probability of formation of a salt bridge between R344 and the Stx1 acidic cluster was 55±25% for the complex with Stx1 S14A compared to 15±10% for the complex with Stx1 S14D. Notably, the IC gate K66-D345 salt bridge has a higher probability of forming in the complex of hDAT SD with Stx1 S14A (60±25%) (Fig. 4D), compared to that of hDAT SD complexed with Stx1 pS14 (17±15%) (Fig. 3D). Thus, the opening of the IC gate and the formation of a translocation pathway filled with water molecules (Fig. 3D) is a process that requires both Stx1 and hDAT N-terminus phosphorylation, a mechanism we believe supports the expression of CDE.

### CDE Occurs in *Drosophila* Brain

*Drosophila* is a powerful model system to determine *in vivo* processes underlying altered DA function due to its conserved mechanisms of DA neurotransmission^56–57^ and its genetic tractability. We utilized the *fmn Drosophila* (DAT^fmn^) background, which lacks expression of full-length *Drosophila* DAT (dDAT) and serves as a functional knock out^58^, in combination with phiC31-based integration to insert a UAS-driven hDAT WT or hDAT SD to express these hDAT constructs specifically in DA neurons, as previously shown^10–11, 59^. We utilized this model system to define the role played by hDAT N-terminus and Stx1 phosphorylation in AMPH-induced DA efflux *ex vivo* and AMPH-associated behaviors^11,55^.

Utilizing isolated fly brains expressing either hDAT WT or hDAT SD, we measured AMPH-induced DA efflux by amperometry. We excised fly brains to maintain intact DA circuitries and record DA efflux from the posterior protocerebral lateral (PPL1) region, a dense cluster of DA neurons that modulate learning^60–61^ (Fig. 5A). As observed *in vitro* (Fig. 2A), AMPH causes DA efflux in brains isolated from hDAT WT flies (Fig. 5B). Inhibition of Stx1 phosphorylation by orally administering CK2i significantly reduced the ability of AMPH to cause DA efflux (Fig. 5B). Notably, in brains isolated from hDAT SD flies, AMPH reveals a CDE (Fig. 5C), as we observed in hDAT SD cells (Fig. 3B). Furthermore, in brains of hDAT SD flies, oral treatment with CK2i restores the ability of AMPH to cause DA efflux (Fig. 5C). These data further support the notion that CDE requires both Stx1 and hDAT phosphorylation in order to occur. To further demonstrate that AMPH unmasks CDE by acting as a hDAT “blocker” in an *ex vivo* preparation, we performed parallel experiments with COC. We applied COC to brain of both hDAT WT and hDAT SD flies. COC promoted a downward deflection of the amperometric trace exclusively in hDAT SD brains, confirming the expression of CDE in hDAT SD but not hDAT WT flies (Fig. 5D). These results demonstrate that CDE is not an *in vitro* epiphenomenon since it occurs in intact brain preparations.

**Figure 5:**
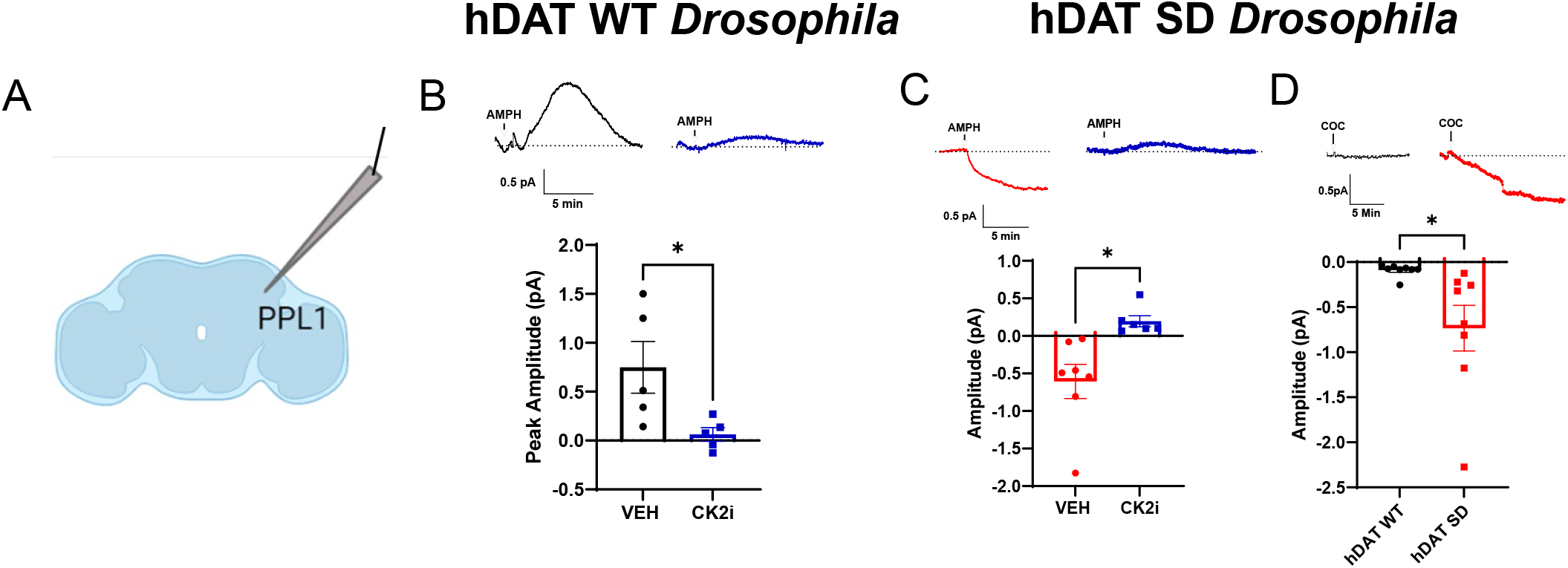
Stx1 and hDAT Phosphorylation Status Regulate DA Efflux in *Drosophila* brains. **A.** Schematic of the amperometric recordings. The amperometric electrode was positioned in the PPL1 region that is enriched in DA neurons. **B.** Representative AMPH-induced amperometric traces (top) recorded from the PPL1 region of hDAT WT expressing *Drosophila* brains from flies who received oral administration of VEH or CK2i (100nM) for 24 hours and quantitative analysis of the peak currents (bottom). CK2i significantly reduced DA efflux in hDAT WT brains (t = 2.505, *n* = 5). **C.** Representative AMPH-induced amperometric traces (top) recorded from hDAT SD expressing *Drosophila* brains from flies who received oral administration of VEH or CK2i (100nM) for 24 hours and quantitative analysis of the peak currents (bottom). CK2i significantly reduced CDE in hDAT SD brains (t = 3.359, *n* = 6-7). **D.** Representative amperometric traces (top) recorded from the PPL1 region of hDAT WT or hDAT SD *Drosophila* brains and quantitative analysis of the peak currents (bottom). Brains were exposed acutely to COC (1 μM). COC reveals a constitutive DA efflux in hDAT SD that is not present in hDAT WT brains (t = 2.528, *n* = 8). Data is presented as mean ± SEM. Student’s unpaired t-test (B); Student’s unpaired t-test with Welch’s correction (C, D).

### CDE and Stx1 Phosphorylation Promote the Expression of Fly Behaviors Associated with AMPH Exposure

To investigate whether CDE, as well as Stx1 phosphorylation, translates to specific behaviors, we first determine their involvement in locomotion, a basic behavior that is regulated by DA across phyla^10, 62–63^. We and others have shown that increases in extracellular DA levels, as a result of DA efflux, promote hyperlocomotion^3, 11–12^. Thus, we determined whether the CDE observed in hDAT SD brains translates to increased locomotion. We compared basal locomotion of hDAT WT and hDAT SD flies. Circadian locomotor rhythmicity over 36 hours was maintained in hDAT SD flies as compared to hDAT WT flies (Fig. 6A, left). However, locomotor activity, as measured by beam crosses over a 24-hour period, was significantly increased in hDAT SD flies (Fig. 6A, right). These results are consistent with the presence of CDE in hDAT SD brains (Fig. 5C), further supporting the ability of DA efflux to regulate specific behaviors. Considering that AMPH inhibits CDE, we next measured locomotion in response to AMPH treatment in hDAT WT and hDAT SD flies (Fig. 6B). While hDAT WT flies exposed to AMPH became hyper-locomotive (Fig. 6B, left), hDAT SD flies exhibit significantly reduced locomotion in response to AMPH (Fig. 6B, right). Thus, inhibition of CDE by AMPH in hDAT SD brains (Fig. 5C) is associated with reduced hyperlocomotion. Next, we determined the role played by Stx1 phosphorylation in AMPH-induced locomotion in hDAT WT and hDAT SD flies. Oral administration of CK2i to hDAT WT flies inhibited the ability of AMPH to cause hyperlocomotion (Fig. 6B, left). No differences were observed between vehicle and CK2i treated flies under basal conditions (Fig. 6B, left). In contrast, in hDAT SD flies, CK2i significantly reduces basal hyperlocomotion with respect to vehicle treatment, supporting the notion that CDE-induced behaviors require Stx1 phosphorylation (Fig. 6B, right). Moreover, CK2i exposure did not further decrease locomotion in AMPH treated hDAT SD flies (Fig. 6B, right).

**Figure 6:**
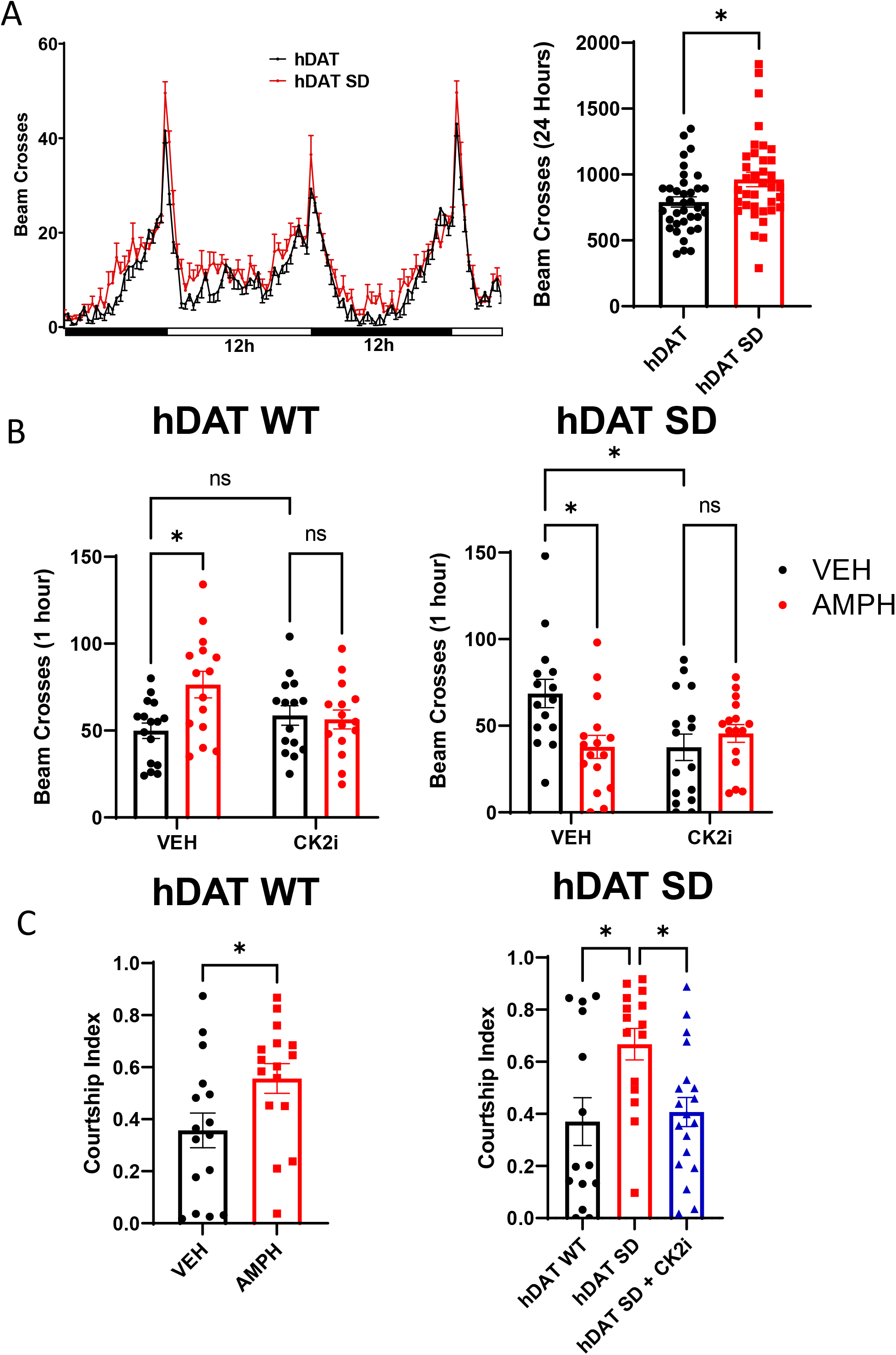
Stx1 Phosphorylation and CDE Regulate AMPH-associated Behaviors in *Drosophila.* **A.** (Left) Locomotor activity was measured by beam crosses over a 36-hour period with a 12:12 light (horizontal white bar)/dark (horizontal black bar) cycle. (Right) Cumulative beam crosses over 24 hours were significantly higher in the hDAT SD compared to the hDAT WT flies (t = 2.556, *n* = 36). **B.** (Left) hDAT WT or (Right) hDAT SD *Drosophila* were orally administered VEH or CK2i (100nm) for 24 hours. Flies were then fed either VEH (black bars) or AMPH (10mM, red bars) and beam crosses were measured over 1 hour. AMPH significantly increased locomotion in hDAT WT flies. This increase was prevented by CK2i treatment (F(1, 57)= 6.037 for effect of interaction, *p* = 0.0157, *n* = 15-16). AMPH and CK2i both significantly decreased locomotion in hDAT SD flies with respect to VEH (F(1, 59) = 7.737 for effect of interaction, *p* = 0.0073, *n* = 15-16). **C.** Courtship index was measured over a 10-minute period. (Left) hDAT WT virgin *Drosophila* males were fed either VEH or AMPH (10mM) for 20 minutes prior to introduction into the 3D chamber to a virgin female hDAT WT *Drosophila.* AMPH significantly increased the courtship index (t = 2.264, *n* = 16). (Right) hDAT WT or hDAT SD virgin *Drosophila* males were orally administered either VEH (hDAT WT and SD) or CK2i (100nM, SD only) for 24 hours and introduced to a virgin female hDAT WT *Drosophila.* hDAT SD flies had a significantly higher courtship index than both hDAT WT and hDAT SD fed CK2i (χ^2^(3) = 8.512, *p* = 0.0142, *n* = 14-19). Data is presented as mean ± SEM. Student’s unpaired t-test (A, C); Two-way ANOVA with Tukey’s multiple comparisons test (B); Kruskal-Wallis test (C).

**Figure 7:**
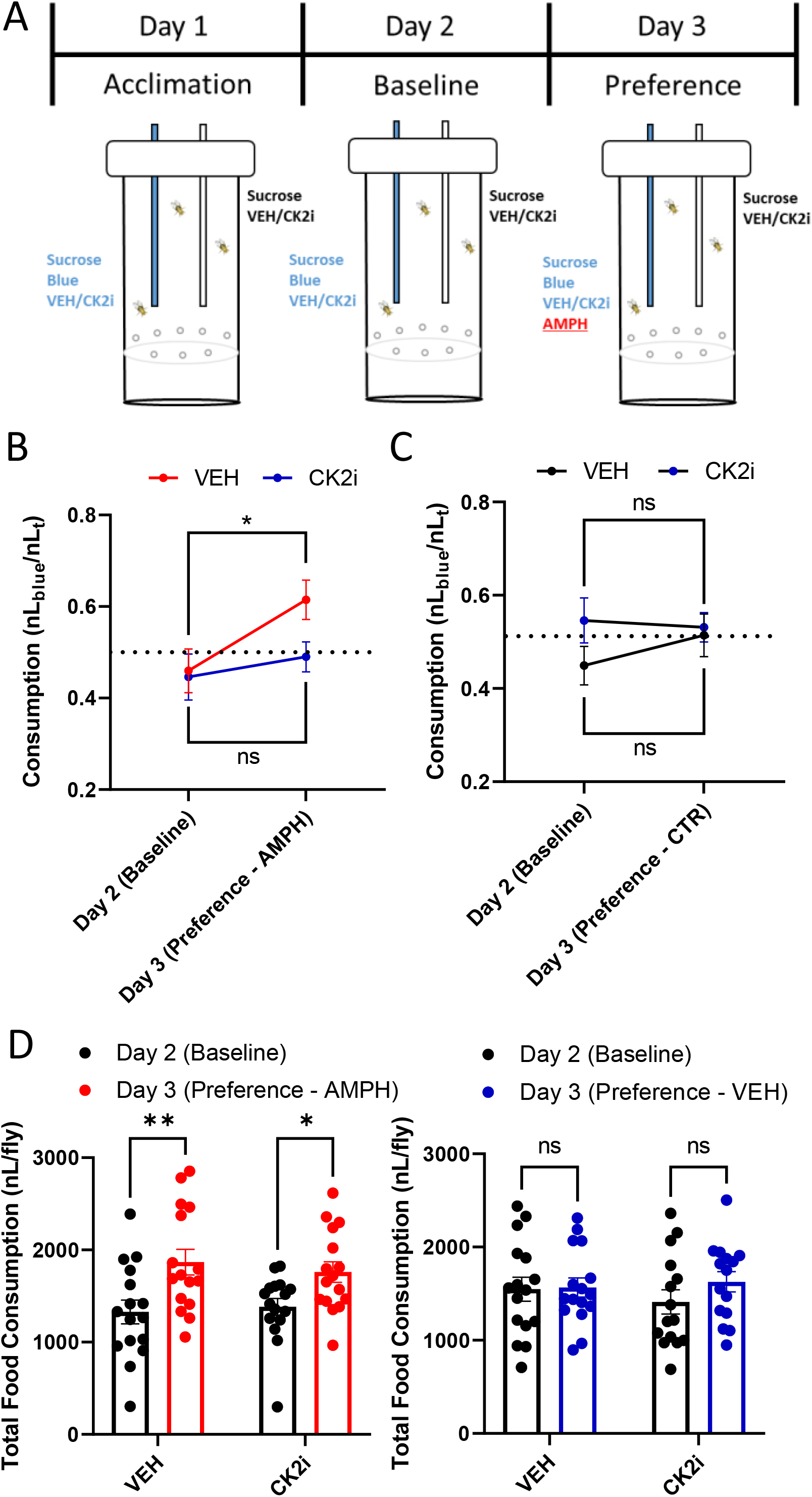
Stx1 Phosphorylation Supports AMPH Preference in *Drosophila.* **A.** Model illustrating a two-choice drug consumption assay developed to measure AMPH preference in flies. The assay is comprised of three 24-hour testing periods: acclimation, baseline, and preference. Flies were acclimated to the two choice apparatus for 24 h (day 1) with food (sucrose) provided in volumetric capillaries. For the duration of the experiment, in half of the apparatuses, sucrose contains CK2i (100nM) and the other half has VEH. In each apparatus one of the two capillaries contains a blue dye in the sucrose. On day 2, food consumption was recorded and the blue capillary food consumption was normalized to the total food consumed (blue capillary/blue+transparent capillaries (nLblue/nLt)). On day 3, AMPH (1 mM) was added to the sucrose of the blue capillaries of half of the apparatuses and VEH to the other half. Food consumption was recorded again. **B.** Flies fed VEH (red line) and given the choice of AMPH (day 3) consumed more from that capillary while flies fed CK2i (100nM, blue line) and given the choice of AMPH consumed an equal amount from both capillaries, the same as the previous day sucrose control (day 2) (F(1, 29) = 5.892 for effect of Day, *p* = 0.0217, *n* = 15-16). **C.** hDAT WT *Drosophila* were treated as above but in the absence of AMPH. These flies demonstrated no preference (day 3) for either of the two capillaries (F(1, 29) = 0.4594 for effect of Day, *p* = 0.5033, *n* = 15-16). **D.** Total food consumption (both clear and blue food combined) was measured for all groups. (Left) hDAT WT flies fed AMPH, regardless of if they received CK2i or VEH, consumed more food on Day 3 than Day 2 (F(1, 30) = 17.53, *p* = 0.0002 for effect of Day; (F(1, 30) = 0.042, *p* = 0.8391 for effect of Drug, *n* = 15-16). (Right) hDAT WT flies that never receive AMPH show no difference in total food consumption (F(1, 29) = 1.100 for effect of Day, *p* = 0.3030; F(1, 29) = 0.0948, p = 0.7603 for effect of Drug, *n* = 15-16). Data is presented as mean ± SEM. Two-way repeated measures ANOVA with Sidak’s multiple comparisons test (B-D).

In *Drosophila,* courting behavior is regulated by DA neurotransmission^64–68^. Therefore, a drug that increases DA signaling, such as AMPH, or a process that increases extracellular DA levels (e.g. CDE) should alter courtship in flies. In these assays, unmated adult male flies harboring hDAT were exposed to vehicle or drug and placed in a 3D-printed chamber. After 10 minutes of acclimation, an adult unmated female fly was introduced and behavior was video recorded for 10 minutes then scored for courtship behavior. Courtship index is calculated as the ratio of time spent demonstrating a courting behavior to total time in the chamber (see methods). Treatment with AMPH significantly increased courtship in hDAT WT flies (Fig. 6C, left; Videos 1-2) underscoring that DA efflux promotes courtship. Consistent with the notion that DA efflux supports courtship, hDAT SD flies exposed to vehicle also demonstrate increased courtship as compared to hDAT WT flies (Fig. 6C, right; Video 3). This hDAT SD phenotype was reversed by treatment with CK2i (Fig. 6C, right; Video 4), as observed for locomotion (Fig. 6B) and CDE (Fig. 5C). However, neither AMPH nor the hDAT SD mutant alter latency to courtship engagement (time it takes the male fly to begin courting the female fly) or the percentage of flies that achieved copulation (Supp. Fig. 7). These data further demonstrate that CDE enhances specific fly behaviors associated with DA neurotransmission and that Stx1 phosphorylation is a key component in this process.

### Stx1 phosphorylation is required for the development of AMPH Preference

Reward is regulated by DA across animal species, ranging from flies to humans^69^. In general terms, the reward component of a natural behavior, particularly feeding, defines how salient a specific behavioral outcome is. Psychostimulants, such as AMPH, work on DA circuits that are designed to respond to natural rewards. In *Drosophila,* we have shown that DA efflux is required for the expression of AMPH preference, a behavior used to quantify the rewarding properties of a psychostimulant^3, 11^. Therefore, we asked whether Stx1 phosphorylation is required for AMPH preference. We adopted a two-capillary feeding assay, developed in our laboratory, for measuring AMPH preference in *Drosophila*^11^. In this assay, adult flies are placed in vials containing two-capillaries; one containing clear sucrose and one containing blue sucrose. Flies are allowed to acclimate for 24 hours (Day 1) then baseline consumption is measured as the ratio of Blue food consumed/Total food consumed (Day 2). A ratio of 0.5 indicates no preference. hDAT WT flies were divided in two groups and treated with either vehicle (VEH), or CK2i, for the entire duration of the experiment. On day 3, either AMPH or control solution (CTR) is added to blue food and consumption in both capillaries is measured for 24 hours (Fig. 7A). Flies fed VEH show a robust AMPH preference (red line) while flies treated with CK2i demonstrate no expression of AMPH preference (Fig. 7B). Furthermore, flies that only receive the VEH or CK2i treatment, without AMPH exposure, demonstrate no preference for clear or blue food (Fig. 7C). These results strongly support role of CK2 activity and Stx1 phosphorylation in the rewarding properties of AMPH.

As previous studies have shown^70^ AMPHs significantly increase food consumption. In our paradigm, we see an increase in total food consumption on day 3 in both the VEH and CK2i groups that were given AMPH (Fig. 7D, left). Of note, is that the AMPH-induced increase in food consumption was not significantly different between VEH and the CK2i group. The groups that did not receive AMPH on day 3 demonstrate no increase in food consumption (Fig. 7D, right).

## Discussion

Stx1 is a member of the SNARE superfamily that plays a critical role in neuronal exocytosis^29^. In addition to its role in vesicular fusion, Stx1 interacts with and regulates the function of transmembrane proteins, including ion channels and transporters^12,32–37^. Mutations and polymorphisms in the Stx1 gene have been implicated in neuropsychiatric disorders such as autism spectrum disorder (ASD)^12, 71–72^ and schizophrenia^73^. Furthermore, changes in post-translational modifications of Stx1 have also been implicated in schizophrenia. A post-mortem study of the prefrontal cortex of patients with schizophrenia demonstrated deficits in both CK2 levels as well as Stx1 pS14^74^, further demonstrating a possible role played by Stx1 and its phosphorylation in the etiology of brain disorders. In addition to ASD and schizophrenia, Stx1 expression levels, function, and phosphorylation have also been implicated in the mechanism of action of AMPH^12, 33, 35, 75^. However, whether and how Stx1 phosphorylation supports specific behaviors associated with AMPH exposure has been largely unexplored. In this study, we defined how Stx1 phosphorylation supports the ability of AMPH to cause DA dysfunction as well as specific behaviors.

We demonstrate that AMPH exposure increases Stx1 pS14 levels as well as hDAT-Stx1 interactions with comparable temporal dynamics, suggesting that Stx1 phosphorylation plays a pivotal role in regulating the association of Stx1 with hDAT (Fig. 1). Consistent with this notion, our molecular modeling shows that Stx1 S14 resides in close proximity to the acidic cluster D15-D19 forming salt bridges with hDAT basic residues R344 and K65/K66 (Fig. 1). Thus, as discussed below, its phosphorylation might cause the structural rearrangements of the Stx1 N-terminus, altering the association of the hDAT with Stx1. We also show that AMPH-induced phosphorylation of Stx1 at S14 is CK2 mediated, since CX-4945 (CK2i) inhibits this phosphorylation event (Fig. 1). The small molecule CX-4945 blocks CK2 activity by competing for the ATP binding site of the kinase and, as a consequence, inhibits phosphorylation of Stx1 at Ser14^50^. It is important to note that CX-4945 is orally bioavailable^76^. Previous data demonstrates that CK2 inhibition and decreased Stx1 pS14 levels selectively impairs the ability of AMPH to cause DA efflux as well as hyperlocomotion^12^. This impairment was also associated with altered hDAT-Stx1 association^12^. The question remains whether and how Stx1 pS14 regulates DA efflux. We first demonstrated that inhibiting Stx1 phosphorylation at S14 either pharmacologically (CK2i) or molecularly (Ser to Ala substitution), significantly impaired DA efflux (Fig. 2). Of note is that cells expressing the pseudo-phosphorylated form of Stx1 (Stx1 S14D) are resistant to the inhibitory action of CK2i, in terms of DA efflux, further reinforcing the fundamental role of CK2-mediated Stx1 phosphorylation in reverse transport of DA. Curiously, the phosphorylation status of the hDAT N-terminus also regulates the levels of Stx1 pS14. Consistent with this notion, preventing phosphorylation of the five hDAT most distal N-terminal Ser by their substitution to Ala (hDAT SA) inhibited the ability of AMPH to increase Stx1 pS14 levels (Fig. 3). In contrast, their pseudophosphorylation (hDAT SD) increased Stx1 pS14 levels under basal conditions (i.e in the absence of AMPH; Supp. Fig. 2). We attribute this hDAT SD-mediated increase in Stx1 pS14 levels to its ability to cause cell membrane depolarization (Supp. Fig. 2). To further support this hypothesis, AMPH, which promotes Stx1 phosphorylation, also causes cell membrane depolarization^15, 20^. Furthermore, there is evidence that CK2 can be activated by membrane depolarization^77^.

CDE is observed upon phosphorylation of Stx1 and hDAT N-terminus. CDE is a phenomenon that has been seen in multiple disease-associated hDAT variations, such as hDAT T356M ^55, 78–79^ and hDAT A559V^53–54, 80^. In the hDAT A559V, CDE is supported, at least in part, by the ability of hDAT A559V to form an aqueous pore^53^. It is important to note that hDAT SD cells, in addition to presenting increased levels of Stx1 pS14, display CDE. Thus, reverse transport of DA can occur in the absence of AMPH once both Stx1 and hDAT N-terminus are phosphorylated (see also below). In hDAT SD cells, CDE was uncovered by its blockade with COC (Fig. 3). hDAT SD-mediated CDE was also blocked by AMPH, a DAT substrate^3^. These data suggest that while hDAT SD cells are capable of DA uptake, AMPH acts like a blocker further pointing to Stx1 phosphorylation as a regulator of DAT function and possibly substrate recognition.

Our MD simulations further elucidated how the phosphorylation state of the hDAT/Stx1 complex modulates their interaction and how this might contribute to reversal of DA transport. We determined that the hDAT-Stx1 interfacial interactions involve salt bridges between Stx1 D15/D17 and hDAT R344, Stx1 R26 and hDAT E56, and Stx1 E38/E39 and hDAT R51 (Fig. 1D). It is important to note that disease-associated hDAT variations resulting in the disruption of these salt bridges (e.g. R26Q in Stx1 or R51W in hDAT) have previously been implicated in altering hDAT-Stx1 interactions and DA efflux^12^. Our 100 ns MD simulation of hDAT SD complexed with Stx1 pS14 shows that pseudophosphorylation of the hDAT N-terminus promotes the breaking of hDAT salt bridges K66-D345 and E428-R445 (Fig. 3D and Supp. Fig. 4), and leads to an inward-facing conformer. In this simulation, there was an influx of water molecules into the vestibule and DA molecules migrate from the cytoplasm space to the vestibule near hDAT E428 (Fig. 3D and Supp. Fig. 4A-C). Considering that hDAT channel-like activity has been shown to support reverse transport of DA^81^, it is possible that Stx1 and hDAT N-terminal phosphorylation promotes CDE by causing, at least in part, the formation of an aqueous pore in hDAT.

The channel-like state of the hDAT SD/Stx1 pS14 complex may also contribute to the paradoxical effects seen in response to AMPH (i.e. AMPH blocks CDE). hDAT SD in the presence of Stx1 pS14 is in a conformation that permits DA to cross the plasma membrane through an aqueous pore, down its electrochemical gradient. Thus, it is possible that application of extracellular AMPH prevents CDE by binding to the hDAT vestibule and occluding the hDAT aqueous pore. The importance of Stx1 phosphorylation in CDE is underscored by the fact that preventing Stx1 phosphorylation (Stx1 S14A) in hDAT SD cells both blocked the expression of CDE and restored, to a minimal level, the ability of AMPH to cause DA efflux (Fig. 4). Thus, Stx1 phosphorylation is essential for the expression of CDE, a functional state of hDAT characterized by reverse transport of DA, which is possibly mediated by the formation of an aqueous pore.

The role played by Stx1 phosphorylation in DA efflux in cells was also observed in isolated brains of *Drosophila.* In hDAT WT fly brains, we demonstrated that CK2 inhibition significantly inhibited DA efflux in response to AMPH (Fig. 5). Notably, as we observed in hDAT SD cells, brains of hDAT SD flies display an AMPH sensitive CDE. Furthermore, in hDAT SD brains, the ability of AMPH to cause reverse transport of DA was restored by pharmacological inhibition of Stx1 phosphorylation (Fig. 5). These data point to the fundamental role played by Stx1 pS14 in CDE, as well as in the ability of AMPH to cause DA efflux, not only in a heterologous expression system, but also in *Drosophila* brains.

To determine the behavioral significance of both CDE and Stx1 phosphorylation, we first determined how hDAT N-terminus and Stx1 phosphorylation alters locomotion in flies. Locomotion is a behavior regulated by extracellular DA^62, 82^. AMPH, by increasing extracellular DA, causes enhanced locomotion in both *Drosophila* and mammals^10–11, 25, 28, 63^. Consistent with the idea that CDE increases extracellular DA levels, hDAT SD flies are hyperlocomotive with respect to hDAT WT flies (Fig. 6). Furthermore, CK2 inhibition by CK2i significantly decreased the hyperlocomotion observed in hDAT SD flies to the level observed in hDAT WT flies. In hDAT SD flies, AMPH also decreases hyperlocomotion, further pointing to CDE as an hDAT function that regulates fly behaviors (Fig. 6). On the other hand, in hDAT WT flies, CK2 inhibition significantly decreased the ability of AMPH to cause hyperlocomotion. These data are consistent with the ability of CK2i to significantly decrease both AMPH-induced DA efflux and CDE.

Another critical DA-associated behavior in flies is courtship^64–67^. DA has been shown to both set and sustain courtship^68^. Upon encountering a potential mate, a male decides whether or not to initiate courtship. Then, at any succeeding moment until copulation, he must decide whether or not to persevere in the courtship that has so far been unsuccessful. Notably, it has been shown that males with high mating drive sustain courtship for minutes, whereas courtship by satiated males, when it occurs, is frequently abandoned^65^. The male’s motivation to court hinges on dopaminergic activity^68^. Consistent with this notion, AMPH, which increases extracellular DA levels, stimulated courtship in hDAT WT flies (Fig. 6). It is important to note that, in flies, an increase in central DA levels can reverse sexual satiety^65^. In line with this observation, hDAT SD flies court significantly more as compared to their WT counterparts (Fig. 6). This increase in sexual motivation is supported by Stx1 phosphorylation and CDE as the significantly enhanced courtship observed by the hDAT SD flies is reduced by CK2i treatment (Fig. 6). These data are notable as AMPHs are commonly used to increase sex motivation in humans^83^ and further demonstrates that the enhanced DA signaling promoted by Stx1 phosphorylation and the hDAT SD mutations have functional behavioral outcomes. While hDAT WT flies, treated with AMPH, and hDAT SD flies showed significantly increased courtship behaviors, these flies failed to have an increase in successful copulation attempts as compared to their respective controls (Supp. Fig. 7). Since only the male flies in this assay were treated with the drug or harbored the hDAT pseudophosphorylation, and there were no changes in successful copulation, we surmise that the receptivity of the female flies to the male’s attempted courtship remained the same regardless of the male’s behaviors. As it has been demonstrated that DA contributes to courtship memory^64^, it should also be considered that hDAT SD flies and flies treated with AMPH might have altered learning in response to denial from a female.

Preference for positive reinforcers is a behavior regulated by DA in animals ranging from humans to flies^69^. Our lab has recently developed a two-choice capillary assay for measuring drug preference in *Drosophila*^11^. Utilizing this system, we addressed the preference for AMPH in hDAT WT flies and examined how inhibiting CK2 and preventing Stx1 phosphorylation impacts AMPH preference. We demonstrated that hDAT WT flies have a robust preference for AMPH and, if the flies are treated with CK2i, this preference is significantly reduced (Fig. 7). There is a well-established link between DA release and DAT availability in reward responses in humans^84–85^. Considering that CK2i inhibits the ability of AMPH to cause DA efflux (Fig. 5), this data further supports a role of DA efflux in AMPH-associated behaviors and of Stx1 phosphorylation in the expression of drug preference. Interestingly, AMPH treatment also significantly increased the total food consumed (food containing VEH plus food containing AMPH), regardless of VEH or CK2i treatments (Fig. 7). Regardless, even with the increased food consumption seen in the group that received AMPH, the group that received CK2i still demonstrated no preference for the drug, indicating that this increased total consumption does not play a role in the preference. Drug reinforcement measures the saliency of a specific drug exposure and is essential for the transition from drug consumption to addiction and dependence^86^. Thus, considering that CK2i (CX-4945) is in Phase II clinical trials for the treatment of cancer^87^, targeting Stx1 phosphorylation for blocking the development of AMPH use disorders might be a new therapeutic opportunity. Taken together, these data elucidate mechanistically how Stx1 phosphorylation supports AMPH-induced DA efflux, CDE, and associated behaviors, including sexual motivation and drug preference.

## Supplemental Figures

**Supp. Figure 1: *In Silico* modeling of the complex between Stx1 and CK2 and how Stx1 Phosphorylation regulate hDAT-Stx1 Interactions. A.** Structural model for the protein complex Stx1 *(green ribbon diagram)* and human CK2 (PDB: 3U4U; *cyan ribbon),* predicted by ClusPro. The Stx1 S14 *(orange,* in VDW format) makes a close contact with the catalytic residue D156 *(red,* VDW format) of the human CK2 consistent with its potential to be phosphorylated. The Stx1 structural model (residues 1-260) was constructed using Robetta based on the rat Stx1 structure in a closed conformation (PDB: 4JEH); the missing loop (residues 1 to 26) in the Stx1 structure was modeled using Robetta. **B-D.** MD snapshots taken from hDAT WT complexed with Stx1 pS14 at (**B**) 5 ns; (**C**) 30 ns and (**D**) 100 ns. *Red* and *blue* VDW spheres represent acidic and basic residues, respectively on both hDAT and Stx1. Residues from hDAT and Stx1 are labeled in bold-face and normal fonts, respectively. During simulations, the phosphorylation of Stx1 S14 led to the disordering of Stx1 N-terminal segment. This disordering promotes the dissociation of the phosphorylated Stx1 S14 and the adjacent sequence of D15-D19 from hDAT R344 and K66, with which they were previously forming salt bridges; disordering of the N-terminal segment is partially stabilized by Stx1 R26 through intermittent formation of a salt bridge with hDAT E56 or E61 (not shown). Similar results were observed in three independent runs. Panel D and Figure 1G were plotted from the same snapshot.

**Supp. Figure 2: Depolarization Causes Stx1 Phosphorylation in hDAT Cells. A.** Representative immunoblot (top) and quantitative analysis (bottom) of hDAT WT or SD cells treated with 10μM AMPH for 10 minutes. AMPH significantly increased Stx1 phosphorylation in hDAT WT but not in hDAT SD cells. hDAT SD led to Stx1 hyperphosphorylation in the absence of AMPH (F(1, 12) = 29.92 for effect of genotype, *p* = 0.0001, *n* = 4). **B.** Membrane potentials measured from either hDAT WT or hDAT SD cells in whole cell current clamp. There was a significant depolarization of the membrane potential in hDAT SD cells (t = 6.273, *n* = 11-13). Data is presented as mean ± SEM. One-way ANOVA with Tukey’s multiple comparisons test (A); Student’s unpaired t-test (B)

**Supp. Figure 3: Surface Expression of hDAT/Stx1 Mutant Combinations. A.** (*top*) Representative immunoblots of the biotinylated fraction from hDAT cells expressing either hDAT WT or variants (hDAT SA, hDAT SD) as well as Stx1 or one of two Stx1 variants (Stx1 S14D, Stx1 S14A). *(bottom)* Quantitative analysis of the immunoblots. The biotinylated fractions were normalized to total expression of either hDAT or Stx1 and expressed as a percent of what observed in hDAT WT + Stx1 cells. There was a significant increase in hDAT surface expression in the hDAT SA + Stx1 S14A group compared to WT hDAT + Stx1 WT, but no other differences between groups were found (F(8, 27) = 5.949, *p* = 0.0002, *n* = 4). There was no change in Stx1 surface expression in any of the groups (F(8, 27) = 1.185, p = 0.2599, *n* = 4). Data is presented as mean ± SEM. One-way ANOVA with Dunnett’s multiple comparisons test (A)

**Supp. Figure 4: Phosphorylation of Stx1 S14 promotes the opening of salt bridge K66-D345 in favor of an IF-like state.** Snapshots were taken from MD simulations of the complex between hDAT SD and Stx1 pS14 at (**A**) 5 ns; (**B**) 20 ns and (**C**) 50 ns. *Red* and *blue* VDW spheres represent acidic and basic residues. Residues from DAT and Stx1 are labeled in bold-face and plain-face, respectively. D_25_RDR_28_ stretch of Stx1, which includes R26, contributes to stabilizing the association of Stx1 with hDAT through its interaction with hDAT E56 (see Fig. 1). The strength of this interaction is enhanced by additional salt bridges between hDAT

S12D and S13D and Stx1 R26. Furthermore, the salt bridge of hDAT D345-K66 broke around 50 ns, leading to an inward-facing intermediate. Similar results were observed in three runs. Panel A and Figure 3D (left panel) were plotted from the same snapshot.

**Supp. Figure 5: PIP2 mediates the association of Stx1 with hDAT.** MD snapshots taken from one run of hDAT WT complexed with Stx1 WT at (**A**) 5 ns; (**B**) 40 ns and (**C**) 100 ns. *Red* and *blue* vDW spheres represent acidic and basic residues. Residues from hDAT and Stx1 are labeled in bold-face and plain-face, respectively. PIP_2_ lipids are colored differently in *licorice* format. During MD simulations, the presence of PIP_2_ lipids coordinate the adjacent binding of aspartates (D15-D19) in Stx1 to hDAT R344 and K66. Note that PIP_2_ may anchor and stabilize the hDAT and Stx1 interaction. The hDAT conformer remained in the *occluded* state during the simulations, with the closure of EC gate R85-D476, and IC gate residues R60-D436, K66-D345, and R445-E428. Similar behavior was observed in two independent runs. Panel C and Figure 1F were plotted from the same snapshot.

**Supp. Figure 6: Stx1 Phosphorylation at S14 requires hDAT N-Terminus phosphorylation to support CDE. A.** Representative traces (top) and quantitative analysis (bottom) of hDAT SA cells expressing either Stx1 WT, Stx1 S14A, or Stx1 S14D. In hDAT SA cells there was no difference in AMPH-induced DA efflux between any of the three Stx1 isoforms (F(2, 10) = 0.9287, *p* = 0.4266, *n* = 4-5). Data is presented as mean ± SEM. One-way ANOVA (A).

**Supp. Figure 7 – AMPH and Stx1 phosphorylation do not regulate either latency to courtship or achieving copulation. A.** hDAT WT male *Drosophila* were fed either VEH or AMPH (10mM) for 20 minutes prior to introduction to the 3D chamber and the female hDAT WT *Drosophila.* Survival analysis was performed on the time it took each male to begin courting the female (latency to court). There was no significant difference between groups in latency to court the female fly (x = 3.426, *p* = 0.0642, *n* = 19). **B.** Total number of hDAT WT flies treated with VEH or AMPH who either achieved or failed to achieve copulation were counted. There was no significant difference between groups. (x = 0.1279, *p* = 0.7206, *n* = 19). **C.** hDAT WT and hDAT SD male *Drosophila* were pretreated with either CK2i (100nM) or VEH for 24 hours and were introduced to a chamber for 10 minutes with a female hDAT WT fly. Survival analysis was performed on the time it took each male to begin courting the female (latency to court). There was no significant difference between groups in latency to court the female fly (x = 1.494, *p* = 0.4738, *n* = 14-19). **D.** Total number of hDAT WT and hDAT SD flies treated with CK2i or VEH who either achieved or failed to achieve copulation were counted. There were no significant differences between any of the three groups (x = 3.145, *p* = 0.2075, *n* = 14-19). Data is presented as mean ± SEM. Mantel-Cox Log-Rank survival analysis (A, C); Chi-square test (B, D)

**Video 1 – Courtship of a hDAT WT male fly fed VEH.** Representative video of a male hDAT WT fly fed VEH courting a female hDAT WT fly. Video is displayed at 5x speed.

**Video 2 – Courtship of a hDAT WT male fly fed AMPH.** Representative video of a male hDAT WT fly fed AMPH courting a female hDAT WT fly. Video is displayed at 5x speed.

**Video 3 – Courtship of a hDAT SD male fly fed VEH.** Representative video of a male hDAT SD fly fed VEH courting a female hDAT WT fly. Video is displayed at 5x speed.

**Video 4 – Courtship of a hDAT SD male fly fed CK2i.** Representative video of a male hDAT SD fly fed CK2i courting a female hDAT WT fly. Video is displayed at 5x speed.

