## Supplementary figures and images for "Syntaxin1 Ser14 Phosphorylation is Required for Non-Vesicular Dopamine Release"

### Supplemental Figure 1

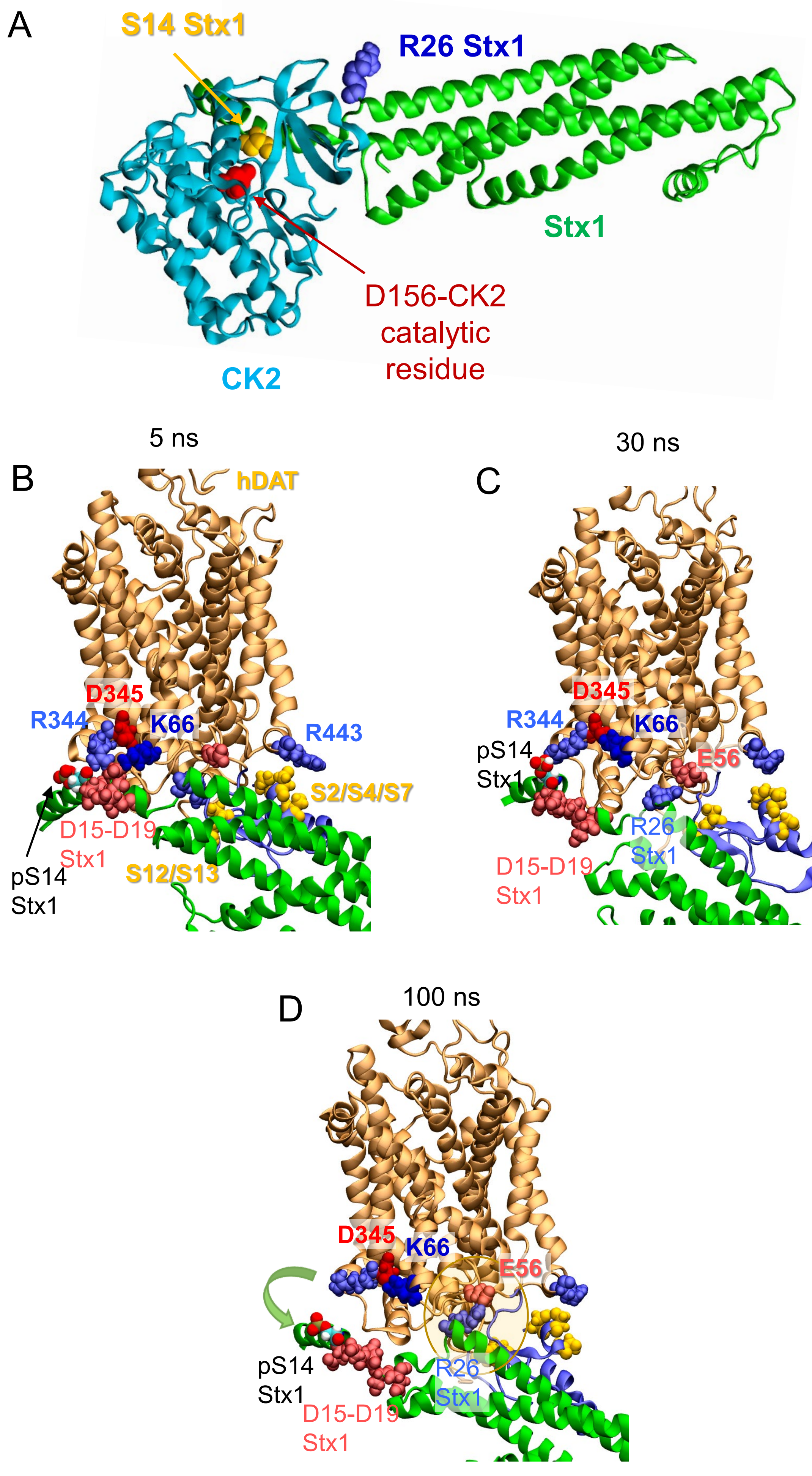

### Supplemental Figure 2

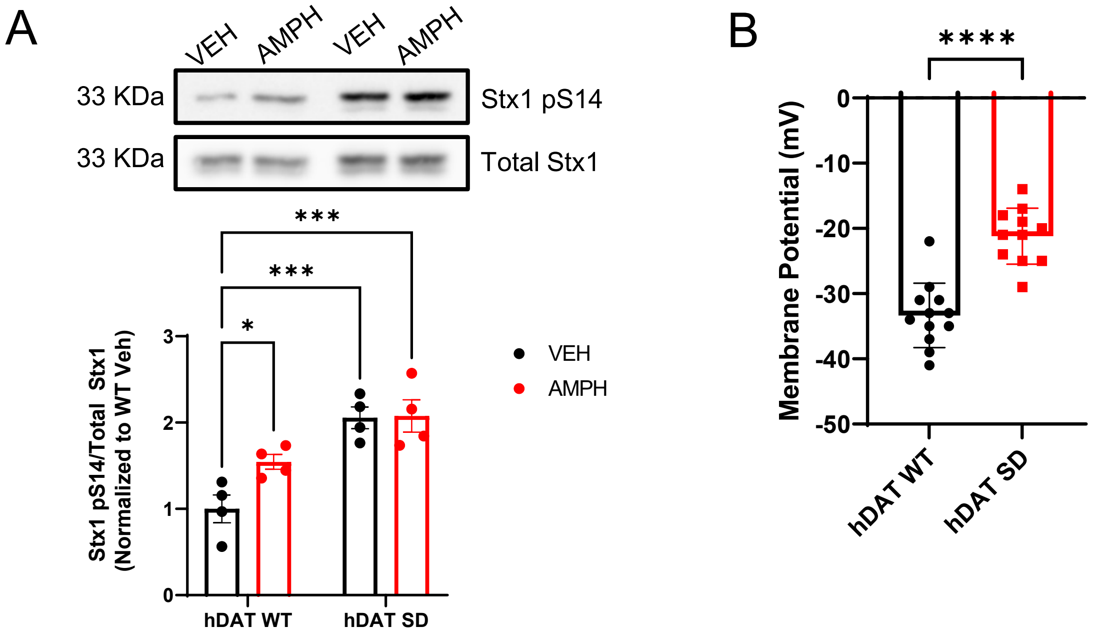

### Supplemental Figure 3

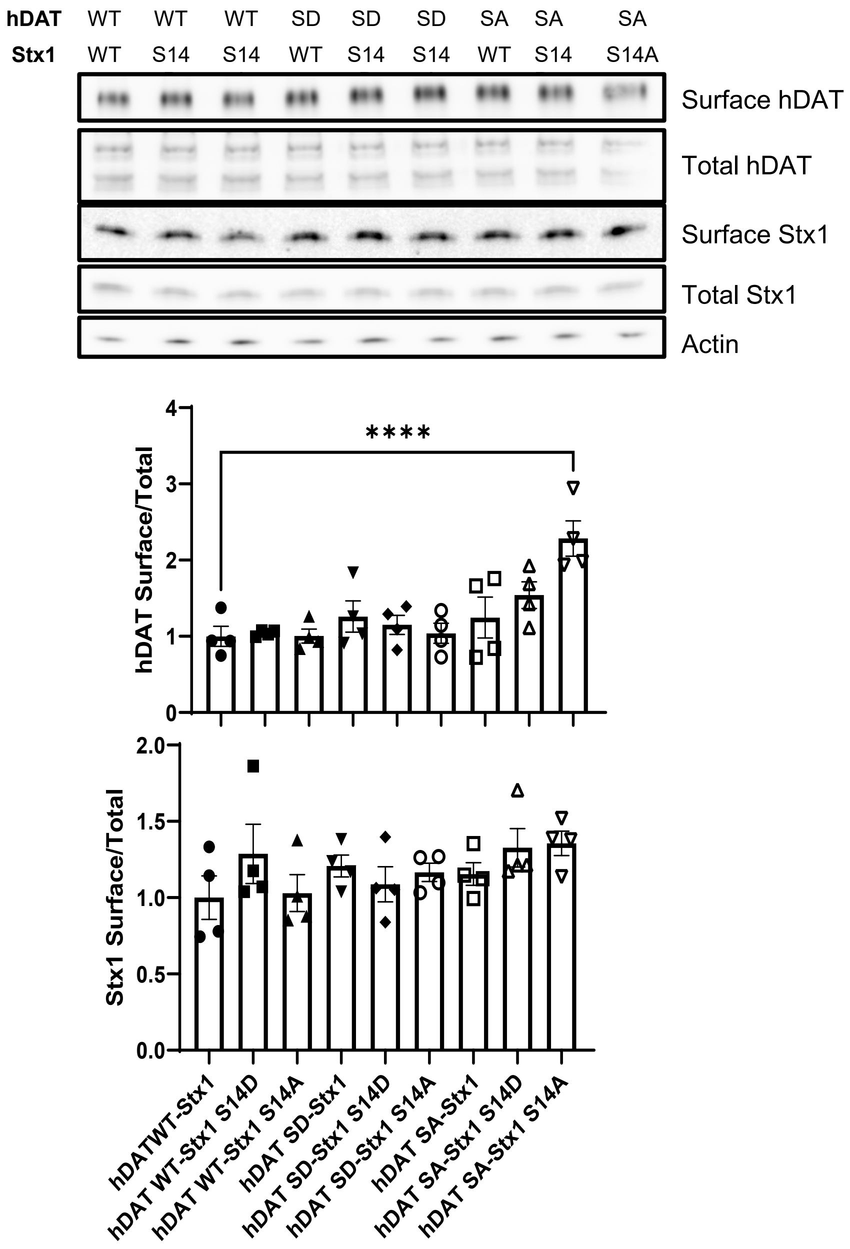

### Supplemental Figure 4

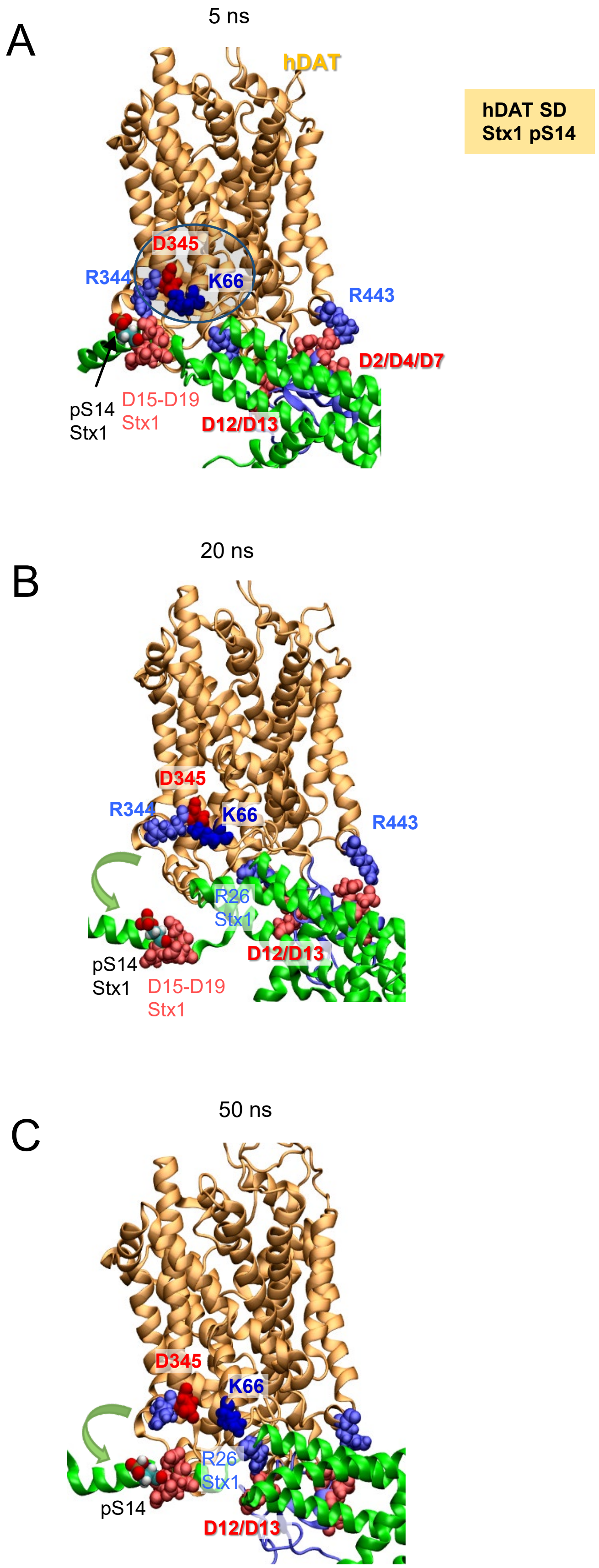

### Supplemental Figure 5

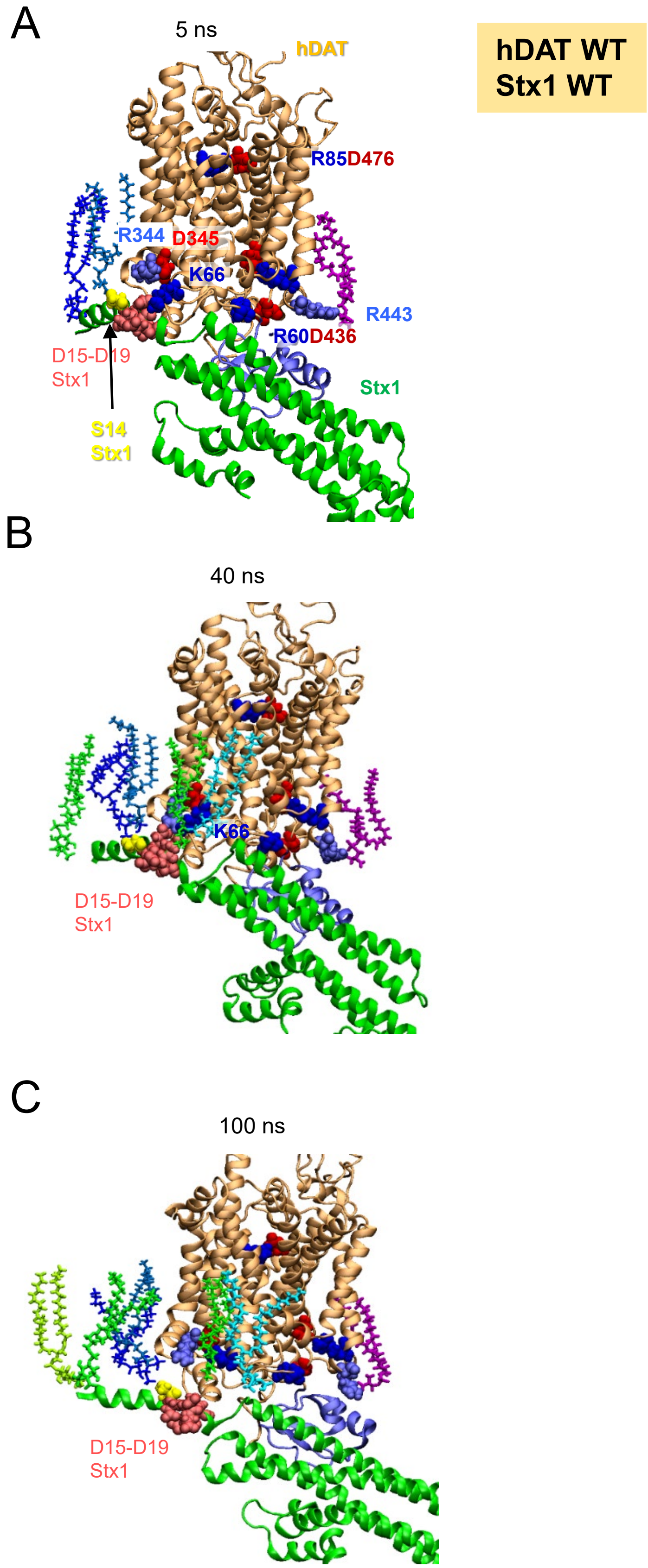

### Supplemental Figure 6

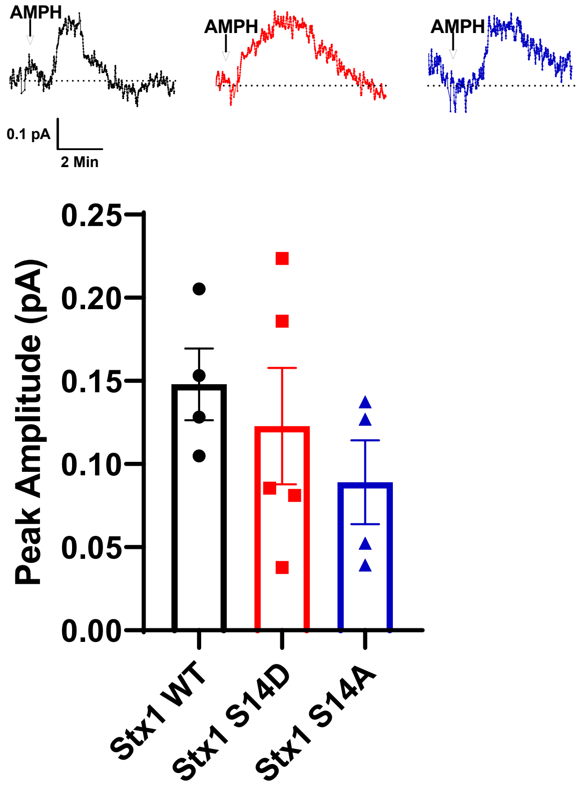
