## Supplemental Figure 7 for "Syntaxin1 Ser14 Phosphorylation is Required for Non-Vesicular Dopamine Release"

A

### hDAT WT

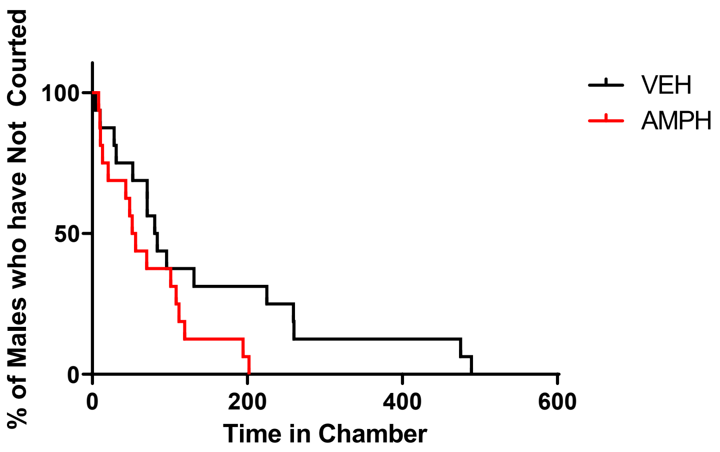


B

| | No Copulation | Achieved Copulation |
|---|---|---|
| hDAT WT + VEH | 13 | 6 |
| hDAT WT + AMPH | 14 | 5 |

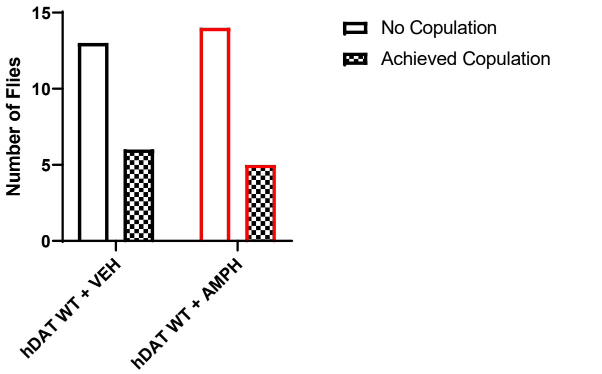


C

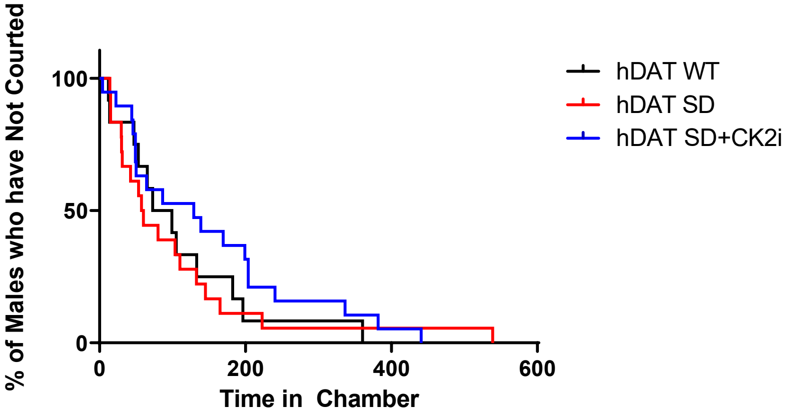


D

| | No Copulation | Achieved Copulation |
|---|---|---|
| hDAT WT + VEH | 13 | 1 |
| hDAT SD + VEH | 13 | 5 |
| hDAT SD + CK2i | 17 | 2 |

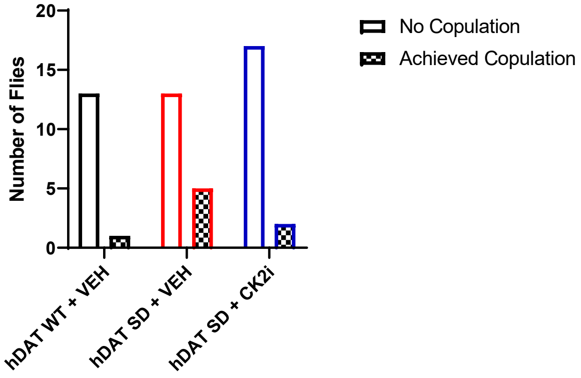
